# The epidermal growth factor (EGF) receptor family network in cardiac hypertrophy and heart failure

**DOI:** 10.64898/2026.05.16.724529

**Authors:** Stephen J Fuller, Susanna TE Cooper, Joshua J Cull, Nikola Adamczyk, Charlotte Tapsell, Riley Pokora, John Spillets, Philip R. Dash, Peter H Sugden, Angela Clerk

## Abstract

The epidermal growth factor receptor (EGFR) network comprises 4 receptors (EGFR, ERBB2, ERBB3, ERBB4) and multiple ligands (EGFR-Ls; for example, AREG/EGF/TGFA that bind to EGFR, EREG/HBEGF that bind to EGFR/ERBB3, and NRG1 that binds to ERBB3/4). NRG1 and ERBB2/ERBB4 are cardioprotective, whilst other EGFR-Ls are implicated in cardiomyocyte hypertrophy. We sought to understand the importance of this network in human heart failure and development of cardiac hypertrophy. By mining data from the Human Protein Atlas, we established that the EGFR family and EGFR-Ls are expressed in different cardiac cells. Furthermore, EGFR-L mRNAs were modulated in failing *vs* non-failing human hearts implicating them in end-stage disease. In rat cardiomyocytes, EGFR-Ls all activated extracellular signal-regulated kinases 1/2 (ERK1/2) and Akt. However, NRG1 preferentially activated Akt whereas AREG/EREG/HBEGF/TGFA preferentially activated ERK1/2. EGF and NRG1 differentially regulated early gene expression with the response to NRG1 having a greater requirement for ERBB2. To determine if the network is involved in developing cardiac hypertrophy, mice were treated with angiotensin II (AngII) without/with afatinib (to inhibit all EGFR family members). Expression of EGFR-L mRNAs was modulated by AngII over 3-7d. At 7d, echocardiography and segmental strain analysis identified greater cardiac dysfunction in hearts from mice treated with afatinib/AngII compared with AngII alone, associated with inhibition of cardiomyocyte hypertrophy and enhanced cardiac fibrosis. In conclusion, the EGFR network plays an important role in cardiomyocyte and cardiac hypertrophy and is dysregulated in human failing hearts. Further insight into the network may enable identification of novel therapies for cardiac disorders.

**Clinical perspectives:**

- The epidermal growth factor receptor (EGFR) network is is potentially important for cardioprotection and cardiac hypertrophy.
- The EGFR family and their ligands (EGFR-Ls) were differentially expressed in human cardiac cells with changes in EGFR-Ls in human heart failure. EGFR-Ls had differential effects on cardiomyocyte signalling/gene expression and were modulated in cardiomyocyte and cardiac hypertrophy. Inhibition of EGFRs in a mouse model of hypertrophy reduced cardiac/cardiomyocyte hypertrophy, enhanced fibrosis and caused cardiac dysfunction.
- The EGFR network is important for cardiomyocyte/cardiac hypertrophy and is dysregulated in human failing hearts. Intervention in this system may offer opportunities to alleviate disease progression.

## Introduction

The biological activity and primary structure of epidermal growth factor (EGF) have been known for over half a century [1]. EGF binds to and activates EGF receptors (EGFRs), the prototypical cell surface tyrosine kinase receptors [2, 3]. Structurally, EGFRs have an extracellular ligand binding domain, a single transmembrane domain and an intracellular C-terminal region containing a tyrosine kinase domain. Mechanistically, EGF binding to the extracellular region activates EGFR dimers with Tyr-phosphorylation of the tyrosine kinase domain. This results in further phosphorylation of other Tyr residues in the intracellular region of the receptors. Proteins are recruited to phospho-Tyr residues and may be subject to further Tyr-phosphorylation, producing a signalosome that initiates intracellular signalling cascades to promote cell proliferation [4, 5]. Two key pathways activated by EGFRs are the prototypical mitogen-activated protein kinases now known as extracellular signal-regulated kinases 1/2 (ERK1/2), and Akt. The EGFR system includes four related receptors [EGFR itself, ERBB2 (HER2 in humans), ERBB3, ERBB4] that are activated by numerous ligands (EGFR-Ls) (**Figure 1A**) [3]. ERBB2/HER2 does not have a functional ligand-binding domain and the tyrosine kinase domain of ERBB3 is inactive but, because they operate as homo- or heterodimers, all EGFR family members have signalling potential. Complexities extend to the EGFR-Ls, some of which have high receptor affinities (e.g. EGF, HBEGF, TGFA) whilst others (e.g. EREG, AREG) have much lower affinities. This can affect signal propagation and, because the EGF system operates locally, acting via autocrine/paracrine networks, the lower affinity ligands have greater potential for diffusion through tissues [6].

**Figure 1.**
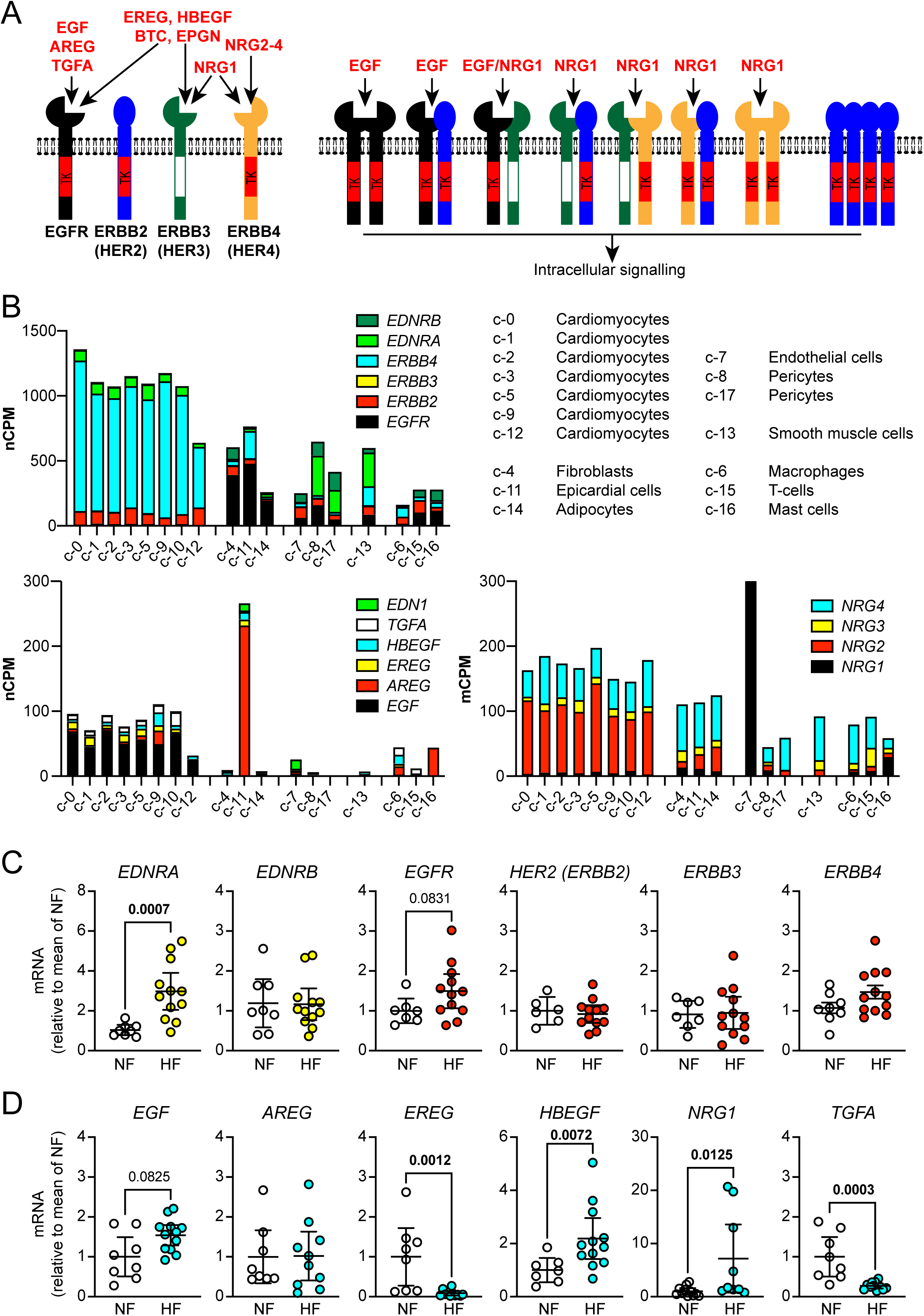
Expression and regulation of the EGFR family and their ligands in human hearts. **A**, Schematic of the EGFR family and the ligands that bind to them. **B**, mRNA expression of receptors (upper panel) and ligands (centre and lower panels) in each of 17 cell types (c-0 to c-17) identified in human heart tissue. Transcript expression of the EGFR family (*EGFR, ERBB2, ERBB3, ERBB4*) is compared with that of the two receptors for endothelin-1 (**EDNRA, EDNRB**). Transcript expression of EGFR family ligands (*EGF, AREG, EREG, HBEGF, NRG1*) is compared with that of endothelin-1 (*EDN1*). Data are from the Human Protein Atlas version 25.0. **C-D**, mRNA expression of endothelin and the EGFR family (**C**) and EGFR family ligands (**D**) in human failing hearts (HF) compared with non-failing hearts (NF). Statistical analysis used Mann-Whitney tests. p values for significant changes (p<0.05) are shown in bold type with values of 0.05-0.1 in standard type.

Activation of EGFRs promotes cell proliferation and cytoprotection which, if unregulated, causes cancer. For example, overexpression of ERBB2 and unregulated activity accounts for 15-20% of all breast cancers [7, 8]. Various therapies target the EGFR system, including antibodies to the extracellular domain and small molecule inhibitors of the intracellular tyrosine kinase [9, 10]. Inhibition of the EGFR family is clearly beneficial for cancer, but causes cardiac dysfunction or heart failure in some patients [11]. This highlights the importance of the EGFR network in the heart.

The heart relies on contractile cardiomyocytes to pump blood carrying oxygen and nutrients to all tissues but, although these cells constitute ∼70% of the volume of the heart, they are only ∼30% of the cell number [12]. Endothelial cells and/or fibroblasts are most numerous, providing, respectively, the capillary network to deliver blood to the heart itself and a robust, but flexible matrix for structure. Other cells include smooth muscle cells in the peri-vasculature. Adult mammalian cardiomyocytes have no significant proliferative capacity [13, 14], but respond to increased workload (e.g. from hypertension) by increasing in size and myofibrillar content (i.e. hypertrophic growth) [15]. This initial adaptation is beneficial and necessary and may be associated with an increase in the capillary network and balanced production of extracellular matrix (i.e. compensated hypertrophy). However, excessive fibrosis and capillary rarefaction can develop, increasing workload whilst reducing oxygen/nutrient availability. This can eventually lead to maladaptation and decompensated hypertrophy [16, 17].

EGFR, ERBB2 and ERBB4 proteins are expressed in human heart samples [18–21]. Single cell proteomics indicates selective expression of ERBB4 in cardiomyocytes, whilst EGFR is expressed in cardiac non-myocytes and ERBB2 is found in all cardiac cells [22]. ERBB2 expression is decreased in human failing hearts, whilst ERBB4 may be up- or downregulated [18, 19, 22]. In addition, phosphoproteomics studies implicate EGFR in cardiac dysfunction associated with hypertrophic cardiomyopathy [21]. These studies indicate that the EGFR network is important in end-stage human heart failure. None of the studies identifies significant expression of ERBB3 in cardiac cells. Egfr, Erbb2 and Erbb4 proteins are readily detected in rat hearts and cardiomyocytes [23–26], and expression of EGFR-Ls is subject to temporal regulation. For example, in neonatal rat cardiomyocytes, the hypertrophic agonist endothelin-1 (ET-1) promotes upregulation of *Areg*, *Ereg* and *Hbegf* mRNAs from 30 min, with maximal expression at 1-2 h returning to basal levels within 4-24 h [27, 28]. A similar pattern is seen with H_2_O_2_ (a physiologically relevant form of oxidative stress) [29]. As in cancer cells, EGF activates ERK1/2 and Akt signalling in cardiomyocytes [30]. These pathways are associated with cardiomyocyte hypertrophy and cytoprotection [31–34], and EGF promotes hypertrophy in rat neonatal cardiomyocytes [30]. HER2/ERBB2 is growth-promoting and cardioprotective in the heart: cardiomyocyte-specific Erbb2 overexpression in mice causes hypertrophic cardiomyopathy, whereas deletion results in dilated cardiomyopathy [35–37]. The latter is attributed to loss of NRG1 signalling via Erbb2/Erbb4 heterodimers. Accordingly, deletion of cardiomyocyte Erbb4 in mice causes dilated cardiomyopathy [38]. NRG1 is considered to be a potential therapeutic option for heart failure [39] and recombinant rhNRG1 is in phase III trials for chronic heart failure (NCT05949801, NCT04468529, NCT03388593).

The EGFR network is clearly important in regulating cardiac function. Detailed understanding of the system is likely to facilitate identification of therapeutic options for amelioration of cardiac disorders whilst also understanding how/why cardiac dysfunction develops from anti-cancer treatments. Here, we determined that mRNAs encoding EGFR-Ls are modulated acutely in neonatal rat cardiomyocytes exposed to EGF or NRG1, in cardiac adaptation to stress in a mouse model of cardiac hypertrophy induced by angiotensin II (AngII), and chronically in human failing hearts. Furthermore, we identified qualitative differences in signalling→gene expression induced by EGF *vs* NRG1 (acting via different EGFR family members) in rat neonatal cardiomyocytes and, in mice *in vivo*, we demonstrated that inhibition of the EGFR family affects cardiac adaptation to AngII. These data confirm the importance of the EGFR network in the heart, extending our understanding of how it operates.

## Methods

### Ethics statement

Human left ventricular samples were from the University of Pittsburgh, U.S.A. (previously reported in [40]). Patients consented to a protocol approved by the University of Pittsburgh Institutional Review Board. Non-failing heart samples were collected with consent being obtained by the local Organ Procurement Organization (OPO), CORE (Center for Organ Recovery and Education) and under University of Pittsburgh CORID #451 (Committee for Oversight of Research and Clinical Training Involving Decedents).

For cardiomyocytes, Sprague-Dawley female rats with 2-4 d litters were purchased from Harlan SeraLab Ltd. UK or Charles River (UK) and housed overnight in the Imperial College Central Biomedical Services facility or the BioResource Unit at University of Reading, respectively. Male C57Bl/6J mice (7 weeks) were purchased from Charles River (U.K.) and imported into the BioResource Unit at University of Reading. All facilities are UK registered with a Home Office certificate of designation. Procedures were performed in accordance with UK regulations and the European Community Directive 86/609/EEC (Imperial College Central Biomedical Services facility) or the European Parliament Directive 2010/63/EU (University of Reading BioResource Unit) for animal experiments. All work was undertaken in accordance with local institutional animal care committee procedures and the U.K. Animals (Scientific Procedures) Act 1986. Neonatal rats were culled by a schedule 1 procedure (cervical dislocation followed by removal of the heart) for which additional Home Office approvals are not required. Experiments with mice were conducted under UK Home Office Project Licences 70/8248, 70/8249 and P8BAB0744.

### mRNA expression of EGFR family and ligands in human cardiac cells

Single cell mRNA data for heart tissue were collected from the Human Protein Atlas (HPA) version 25.0 (release date: 11 November 2025) [41]. Cluster data from GSE183852 [42] (https://www.proteinatlas.org/humanproteome/single+cell/single+cell+type/data#cluster_data) were mined for expression of the EGFR family, EGFR-Ls, endothelin receptors and ET-1 in each of the different cardiac cell types (download date: 23 April 2026). Data are presented as normalised expression (nCPM).

### Preparation and treatment of rat neonatal cardiomyocytes and cardiac non-myocytes

Neonatal female rats with 2-4 d litters were housed in Allentown NexGen Rat 1800 cages with raised wire bar lid, rear shelf, and enrichment loft (58.8cm × 28.6cm × 41.2cm; 1800cm^2^ solid flooring). Cages were supplied with bedding (Eco-Pure Aspen Chips 4, 5 × 5 × 1 mm) plus additional enrichment (nesting material, tunnel, Allentown enrichment loft, aspen chew blocks). Animals were provided *ad libitum* with filtered water via a non-automated bottle watering system and food (irradiated 2018 Global 18%; Envigo) with enrichment (millet and bacon bits). Housing was in an environment with a 12:12 h light/dark cycle, room temperature of 20 ± 1°C and 55 ± 10% humidity.

Neonatal rat ventricular myocytes were prepared and cultured from 3–4 d Sprague–Dawley rats (Charles River) as previously described [43]. Briefly, neonatal rats were culled, and the ventricles dissected and cleaned. Cardiomyocytes were dissociated by serial digestion at 37°C using 0.44 mg/ml (6800 U) Worthington Type II collagenase (from Lonza) and 0.6 mg/ml pancreatin (Sigma-Aldrich, cat. No. P3292). Cells were collected by centrifugation (5 min, 60 × g) and resuspended in maintenance medium [Dulbecco’s modified Eagle’s medium (DMEM) and medium 199 (4:1 (v/v) ratio), 100 U/ml penicillin and streptomycin] containing 15% (v/v) foetal calf serum (FCS; Life Technologies). Non-adherent cardiomyocytes were collected following pre-plating onto plastic tissue culture dishes (30 min) to remove cardiac non-myocytes, allocating cells equivalent of 10 hearts per 60 mm dish. Viable cells were counted by Trypan Blue (Sigma-Aldrich) exclusion using a haemocytometer. Cardiomyocytes were plated onto 60 mm Primaria dishes pre-coated with sterile 1% (w/v) gelatin (Sigma–Aldrich) (4 × 10^6^ cells/dish) and cultured for 18 h. Cardiac non-myocytes were cultured from the pre-plates generated during the preparation of cardiomyocytes. Cells were cultured in maintenance medium containing 5% (v/v) FCS (7 d).

Cells were incubated in maintenance medium for 24 h prior to experimentation, then treated with ligands at the concentrations and for the times indicated. EGF and ET-1 were from Sigma-Aldrich. AREG, HBEGF, NRG1 and TGFA were from R&D Systems. For inhibitor studies, cells were pre-incubated with 0.1% DMSO, 1 µM afatinib or 1 µM tucatinib in DMSO (Selleck Chemicals) for 15 min before addition of EGF or NRG1. Peptide growth factors were stored as 100 µg/ml stock solutions prepared according to the manufacturer’s instructions. ET-1 was stored as a 1 µM stock solution in purified water. Afatinib and tucatinib were prepared as 1 mM stock solutions in DMSO and stored at −20°C.

### Western blotting

Cardiomyocytes were washed with ice-cold PBS, scraped into 150 µl of extraction buffer [20 mM β-glycerophosphate pH 7.5, 50 mM NaF, 2 mM EDTA, 1% (v/v) Triton X-100, 5 mM dithiothreitol, 10 mM benzamidine, 0.2 mM leupeptin, 0.01 mM trans-epoxy succinyl-l-leucylamido-(4-guanidino)butane, 0.3 mM phenylmethylsulphonyl fluoride, 4 µM microcystin]. Samples were vortexed in Eppendorf safe-lock tubes and extracted on ice (10 min). Insoluble material was pelleted by centrifugation (10 000 × g, 10 min, 4°C). The supernatants were collected, a sample was taken for protein assay and the remainder boiled with 0.33 vol sample buffer (300 mM Tris–HCl pH 6.8, 10% (w/v) SDS, 13% (v/v) glycerol, 130 mM dithiothreitol, 0.2% (w/v) bromophenol blue). Protein concentrations were determined by Bio-Rad Bradford assay using BSA standards. Proteins were separated by SDS–polyacrylamide gel electrophoresis on 10% (w/v) polyacrylamide resolving gels with 6% stacking gels, then transferred to nitrocellulose using a Bio-Rad semi-dry transfer cell (10 V, 60 min). Non-specific binding sites were blocked with 5% (w/v) non-fat milk powder in Tris-buffered saline (TBST: 20 mM Tris–HCl pH 7.5, 137 mM NaCl containing 0.1% (v/v) Tween 20) (15 min). Blots were incubated with primary antibodies (1/1000) in TBST containing 5% (w/v) BSA (overnight, 4°C), washed with TBST (3 × 5 min, 21°C), incubated with horseradish peroxidase-conjugated secondary antibodies (1/5000 dilution in TBST containing 1% (w/v) non-fat milk powder, 60 min, 21°C) and washed again in TBST (3 × 5 min, 21°C). See **Supplementary Table S1** for details of antibodies. For time-course and concentration-dependency experiments, bands were detected by enhanced chemiluminescence (Santa Cruz Biotechnology) using Hyperfilm MP with quantification by scanning densitometry (ImageMaster 1D Gel Analysis v3.0quant software; GE Healthcare, now Cytiva). For studies with tucatinib, bands were detected using ECL Prime Western Blotting detection reagents (Cytiva) and visualised using an ImageQuant LAS4000 system (GE Healthcare). ImageQuant TL 8.1 software (GE Healthcare) was used for densitometric analysis. Raw values for phosphorylated kinases were normalised to the total kinase. Values for all samples were normalised to the mean of the controls. Original, full-size images of the scans of X-ray films with original labelling and digital images are provided in the Supplementary information (**Supplementary Source Figures SF1-SF2**).

### RNA preparation and qPCR

Human left ventricular or mouse heart powders (10-20 mg) were weighed into Eppendorf Safe-Lock tubes on dry ice. RNA Bee (1 ml; AMS Biotechnology Ltd) was added as the tube was brought to 4°C. Cardiomyocytes were scraped into 0.5 ml RNA Bee per 60 mm dish, transferred to Eppendorf Safe-Lock tubes, and the dish was washed with 0.5 ml RNA Bee which was added to the same tube. Samples were homogenised in the tubes on ice using a plastic pestle. RNA was prepared according to the manufacturer’s instructions and as described in [43]. RNA was dissolved in nuclease-free water and the purity assessed from the A_260_/A_280_ measured using an Implen NanoPhotometer. Samples with values of 1.8–2.0 were used and concentrations were determined from the A_260_. RNA was stored at −80°C. Total RNA was used to prepare cDNA using High Capacity cDNA Reverse Transcription Kits with random primers (Applied Biosystems) in a final volume of 20 µl according to the manufacturer’s instructions. cDNAs were diluted 1:1 with nuclease-free water and stored at −20°C. qPCR was performed with a StepOnePlus Real-Time PCR system (ThermoFisher Scientific) using 1 µl of the cDNA. Optical 96-well reaction plates were used with iTaq Universal SYBR Green Supermix (Bio-Rad Laboratories Inc.) according to the manufacturer’s instructions. See **Supplementary Table S2** for primer sequences. Results were normalized to *Gapdh*, and relative quantification was obtained using the ΔCt (threshold cycle) method: relative expression was calculated as 2^−ΔΔCt^, and normalised to the mean of the control hearts (patients) or control cardiomyocyte samples.

### RNA-sequencing (RNAseq)

Total RNA (A_260/280_ ratios 1.9-2.0) was diluted and provided to Novogene (UK) Ltd for quality control assessment. RNA was prepared and sequenced according to their protocols. In brief, mRNA was purified using poly-T oligo-attached magnetic beads and fragmented. First strand cDNA was synthesized using random hexamer primers, followed by second strand cDNA synthesis using dTTP for non-directional libraries [44]. Samples were subjected to end repair, A-tailing, adapter ligation, size selection, amplification, and purification. The library was checked with Qubit and real-time PCR for quantification, and bioanalyzer for size distribution detection. Quantified libraries were sequenced on Illumina platforms.

Bioinformatics was performed by Novogene (UK). Raw data (raw reads) in FASTQ format was processed through in-house perl scripts. Clean data (clean reads) were obtained by removing low quality reads and reads containing adapter or poly-N from raw data. Q20, Q30 and GC content were calculated. All downstream analyses were based on the cleaned data with high quality. To map reads to the reference genome, the reference genome and gene model annotation files were downloaded from the genome website directly. The index of the reference genome was built using Hisat2 v2.0.5 [45] and paired-end clean reads were aligned to the reference genome using Hisat2 v2.0.5. The mapped reads of each sample were assembled by StringTie [46] (v1.3.3b) in a reference-based approach. FeatureCounts [47] (v1.5.0-p3) was used to count the reads numbers mapped to each gene. The FPKM (Fragments Per Kilobase of transcript sequence per Millions base pairs sequenced) for each gene was calculated based on the length of the gene and reads count mapped to this gene. Differential expression analysis of two conditions/groups (two biological replicates per condition) was performed using the DESeq2R package (1.20.0) that provides statistical routines for determining differential expression in digital gene expression data using a model based on the negative binomial distribution [48, 49]. The resulting P-values were adjusted using the Benjamini and Hochberg’s approach for controlling the false discovery rate. Genes with an adjusted p-value <=0.05 found by DESeq2 were assigned as differentially expressed. The clusterProfiler R package was used to test the statistical enrichment of differentially expressed genes in KEGG pathways [50]. The data are available from ArrayExpress (E-MTAB-15921) and can be accessed at https://www.ebi.ac.uk/biostudies/ArrayExpress/studies/E-MTAB-15921?key=d97f807e-4628-49f5-b6b2-12cd4334b3d9.

For downstream in-house analysis, differentially expressed transcripts for EGF, NRG1 and tucatinib *vs* controls, tucatinib/EGF *vs* EGF and tucatinib/NRG1 *vs* NRG1 were assembled into a single list (3309 genes). Genes were filtered according to FPKM >0.5 for the mean value of any condition and those with fold-change <1.25 for EGF, NRG1 or tucatinib *vs* controls (1683 genes; **Supplementary Spreadsheet SS1**). ClustVis [51] was used with these genes for Principal Components Analysis (PCA) of samples and heatmap generation.

### Treatment of mice with AngII and afatinib

The experimental unit was the individual mouse and a total of 30 mice are reported in this study (n=9 for AngII and gene expression at 3 d; n=21 for AngII with afatinib). Mice were housed in Tecniplast IVC cages (total area 512 cm^2^; maximum five mice per cage) supplied with aspen sawdust bedding, sizzle nesting, cardboard tunnels and housing plus additional enrichment (e.g. chew sticks and millet). They were provided with water and food (SDS Rm3 pelleted food) *ad libitum*, with a 12:12 h light/dark cycle and room temperature of 21°C. Mice were allowed to acclimatise for 7 d prior to experiments. All animals were checked at least once a day by a trained, competent person and licence holders informed of any welfare issues, with consultation with a Named Veterinary Surgeon when necessary. Mice were allocated to groups on a random basis and none was excluded after randomisation. Individuals conducting the studies were not blinded to experimental conditions for welfare monitoring purposes. Animals were checked daily and mice undergoing procedures were monitored using a score sheet. Weights were taken before and at the end of the experiment (**Supplementary Table S3**). All mice are routinely culled if they reach a predefined endpoint agreed with the Named Veterinary Surgeon but no mice reached this point in the experiments included in this study. No other exclusion criteria were set *a priori* for the experimental process.

Alzet osmotic minipumps (1007D; supplied by Charles River, UK) were used to deliver acidified PBS vehicle or 0.8 mg/kg/d AngII in acidified PBS. For experiments with afatinib, all mice were implanted with a second minipump filled with DMSO/PEG mix [50% (v/v) dimethyl sulphoxide (DMSO), 20% (v/v) polyethylene glycol 400, 5% (v/v) propylene glycol, 0.5% (v/v) Tween 80] or 4.5 mg/kg/d afatinib (Selleck Chemicals) in DMSO/PEG mix. This was based on the recommended dose of afatinib to inhibit EGFR in patients which is up to 50 mg/d p.o. Up to 1400 mg/d could therefore be necessary to inhibit ERBB2 (the IC_50_ values for EGFR and HER2 are 0.5 and 14-16 nM, respectively) [52, 53]. Assuming an average patient weight of 80 kg and bioavailability of ∼25-30% suggests a daily dose of 4.4 - 5.2 mg/kg/d should inhibit all EGFR family members and a dose at the lower end of this spectrum was selected. Minipumps were filled according to the manufacturer’s instructions in a laminar flow hood using sterile technique. and were incubated overnight in sterile PBS (37°C). Mice were given 0.05 mg/kg (s.c.) buprenorphine (Ceva Animal Health Ltd.) for analgesia. Minipumps were implanted under continuous inhalation anaesthesia [using isoflurane (induction at 5%, maintenance at 2–2.5%) mixed with 2 l/min O_2_] via an incision in the mid-scapular region and a pocket created in the left flank region of the mouse. Minipumps were implanted portal first into this pocket. Wound closure used a simple interrupted suture with polypropylene 4-0 thread (Prolene, Ethicon). Mice were allowed to recover and returned to their home cage. At the end of the experiment, mice were culled by CO_2_ inhalation followed by cervical dislocation. Hearts were excised quickly, washed in PBS, dried and fixed with 10% formalin or snap-frozen in liquid N_2_. Fozen hearts were ground to powder under liquid N_2_ and stored at −80°C.

### Histological staining

Histological staining and analysis were performed as previously described, assessing general morphology by haematoxylin and eosin (H&E) and fibrosis by Masson’s trichrome and picrosirius red (PSR). Briefly, hearts were fixed with 10% formalin. Following immersion in 70% ethanol, hearts were embedded in paraffin, sectioned at 10 μm and stained using kits for H&E (Sigma) or Masson’s trichrome (Polysciences). For PSR, sections were submerged in Weigert’s hematoxylin, washed, stained in picrosirius red (1 g/L Sirus red in saturated aqueous picric acid, 60 min), and differentiated in 0.5% acetic acid. Images of heart sections were captured and stored digitally using a Nikon slide scanner. For analysis of myocyte cross-sectional area, cells stained by H&E within the LV (excluding endocardial regions) were chosen at random and outline traced using NDP.view2 software (Hamamatsu). Only cells with a single nucleus were included in the analysis. For assessment of interstitial and perivascular fibrosis at the percentage level, 20X Massons-trichrome and PSR images of the entire LV were exported and the collagen fraction calculated as the ratio between the sum of the total area of fibrosis (red colour) to the sum of the total tissue area (including the myocyte area) for the entire image using ImageJ. For perivascular fibrosis, the degree of fibrosis was also scored (0, negligible; 1, limited fibrosis; 2, extensive fibrosis around a single vessel or clear fibrosis around more than one vessel; 3, extensive fibrosis around more than one vessel; 4, extensive fibrosis around more than one vessel, permeating into the myocardium). All histological and data analysis was performed by independent assessors blinded to treatment groups. Uncropped images are in the Supplementary information (**Supplementary Source Figure SF3**).

### Echocardiography

Echocardiography was performed using a Vevo 2100^TM^ (Fujifilm Visualsonics) with a 38 MHz MS400 transducer. Baseline echocardiograms were collected before minipump implantation and at 3 and 7 days post-minipump implantation. Anaesthesia was induced using vaporised 5% isoflurane with 1 l/min oxygen; maintenance of anaesthesia used 1.5% isoflurane in 1 l/min oxygen delivered with a nose cone. Mice were placed on a heated physiological monitoring stage in a supine position, and heart rate, respiration rate and body temperature were monitored. Chest fur was removed using an electric razor and hair removal gel, pre-warmed ultrasound gel was applied to the chest, and the transducer lowered into the gel until a clear image was obtained. Imaging was completed within 10-15 min. Mice were recovered singly and transferred to the home cage. Left ventricular (LV) cardiac function and dimensions were assessed from long axis B-mode images. Original images are provided in the Supplementary information (**Supplementary Source Figure SF4**). Speckle-tracking (strain) analysis was performed using VevoStrain software (Visualsonics). Analysis was performed across two cardiac cycles starting immediately after a breath had been taken, with the associated movement of the heart. Time-to-peak analysis was performed to obtain data for different segments of the LV endocardial wall. Segmental data are provided in **Supplementary Spreadsheet SS2**.

### Statistical analysis and presentation of data

Data analysis used Microsoft Excel with GraphPad Prism 10.6.1 for statistical analysis and data presentation. Outliers were excluded from the analysis using a Grubb’s outlier test (decided *a priori*) removing a maximum of a single outlier per condition. Statistical testing used Mann-Whitney tests for comparisons of gene expression in failing *vs* non-failing hearts and effects of AngII on EGFR ligand expression. Time-course studies in cardiomyocytes and histology/gene expression in hearts from mice treated with vehicle, AngII or afatinib/AngII were assessed using Kruskal-Wallis tests with Dunn’s multiple comparisons post-hoc tests. Echocardiography data were analysed using two-way paired (according to time) ANOVA with Tukey’s multiple comparison post-test. Effects of afatinib or tucatinib on cardiomyocyte signalling and gene expression used two-way ANOVA with Tukey’s multiple comparison post-test. Specific p values are provided with significance levels of p<0.05 in bold type.

Images from proprietary software were exported as .jpeg files and cropped for presentation with Adobe Photoshop CC maintaining the original relative proportions. Original images are in **Supplementary Source Files SF1-SF4**.

## Results

### The EGFR family and their ligands in cardiomyocytes and the heart

Of the four EGFR family members (**Figure 1A**), Erbb2 and Erbb4 proteins are expressed in cardiomyocytes (human, rat neonatal/adult), whilst Egfr protein is primarily expressed in cardiac fibroblasts (neonatal rat and human) and smooth muscle cells (human) [18–26]. However, there are different populations of cardiomyocytes that can be defined using single cell transcriptomics, in addition to cardiac fibroblasts and other cardiac non-myocytes, and the expression profiles in each of these cell types is unknown. We therefore mined single cell mRNA data from the Human Protein Atlas (HPA) to identify the receptors expressed in different cardiac cells, comparing the EGFR family with endothelin receptors that have an established role in cardiac hypertrophy [54]. All EGFR family members were expressed in cardiomyocytes but to varying degrees (*ERBB4*>>>*ERBB2*>>*EGFR*>>*ERBB3*) (**Figure 1B, upper panel**; **Supplementary Table S4**). *ERBB4* mRNA was expressed at ∼10-fold higher levels than *ERBB2* which was expressed at a similar level as *EDNRA*. *ERBB3* mRNA was expressed at particularly low levels and was undetectable in some populations. *EGFR* mRNA was expressed particularly in fibroblasts, epicardial cells and adipocytes, whereas mRNAs for endothelin receptors were expressed at higher levels in endothelial cells, pericytes and smooth muscle cells. There was limited expression of *ERBB4* mRNA in cardiac non-myocytes compared with cardiomyocytes. These profiles provide detailed information of the variation in mRNA profiles for the EGFR family in different populations of cardiac cells.

EGFR-Ls most likely act in a local paracrine/autocrine manner [6]. In such situations, constitutive expression can maintain homeostasis, whilst modulated expression signals adaptation to a changing environment. Single cell HPA mRNA profiles indicated that expression of EGFR-Ls in different cardiac cell types varied at baseline, with higher *EGF* expression in cardiomyocytes and *AREG* expression in epicardial cells (**Figure 1B**, lower left panel**; Supplementary Table S6**). Of the neuregulins, *NRG4* was expressed in most cardiac cells with almost exclusive expression of *NRG1* in endothelial cells and *NRG2* in cardiomyocytes (**Figure 1B**, lower right panel**; Supplementary Table S6**). These data point to the likely importance of neuregulin signalling in maintaining homeostasis with autocrine/paracrine signalling to cardiomyocytes, whilst paracrine EGF signalling from cardiomyocytes to cardiac non-myocytes may also be a significant factor. As a low affinity ligand, *AREG* has potential for longer distance signalling [6] which may be significant for epicardial cells.

To assess possible involvement of the EGFR family and EGFR-Ls in heart failure, we used qPCR to determine expression in RNA samples from the left ventricle (LV) of non-failing and failing hearts. Transcripts for the *EDNRA* receptor were upregulated in samples from failing hearts as expected [55]. Expression of mRNAs for the EGFRs did not change significantly, although there was a suggestion that *EGFR* and *ERBB4* expression may increase in some patients (**Figure 1C**). Of the EGFR-Ls, *HBEGF* and *NRG1* mRNAs were significantly upregulated, whilst *EREG* and *TGFA* mRNAs were downregulated in failing relative to non-failing hearts (**Figure 1D**). Thus, changing expression of EGFR-Ls may be important in end-stage disease.

### EGF family receptor signalling to gene expression in rat neonatal cardiomyocytes

Our previous studies highlighted temporal changes in gene expression induced in cardiomyocytes by hypertrophic agonists such as ET-1, including robust and transient expression of EGFR-Ls [27, 28]. For those studies, we used primary rat neonatal cardiomyocytes (from P3-4 neonates for limited proliferative potential) as the most suitable cell model for studying cardiomyocytes signalling to gene expression. Adult cardiomyocytes undergo apoptosis without additional supplements such as insulin (see, for example, [56]) that activate Akt signalling [31, 32, 57], precluding study of this pathway. We therefore used rat neonatal cardiac cells to determine the effects of EGFR-Ls on signalling and gene expression.

We first assessed the relative distribution of EGFR family mRNAs in rat neonatal cardiac cells using qPCR. As expected, cardiomyocytes expressed high levels of cardiomyocyte markers (*Myh6*, *Myh7*, *Nppb*) (**Figure 2A**, left panel). Cardiac non-myocytes had high levels of the smooth muscle cell marker *Tagln* (*Sm-22a*; [58]), with expression of receptors for angiotensin II (*Agtr1a*) and the fibroblast marker *Ddr2* [59] (**Figure 2A**, centre panel). Thus, although this is a mixed culture, smooth muscle cells are the dominant cell type. Consistent with HPA data (**Supplementary Table S4**), *Erbb4* mRNA expression was relatively greater in cardiomyocytes, *Erbb3* mRNA was higher in cardiac non myocytes and *Erbb2* mRNA was approximately equally distributed (**Figure 2A**). *Egfr* mRNA was approximately equally expressed in cardiac myocytes and non-myocytes (**Figure 2A**, right panel), so there may be higher expression in neonatal cardiomyocytes than in adult cells, but it is possible to compare signalling from EGFRs, ERBB2 and ERBB4 in these cells.

**Figure 2.**
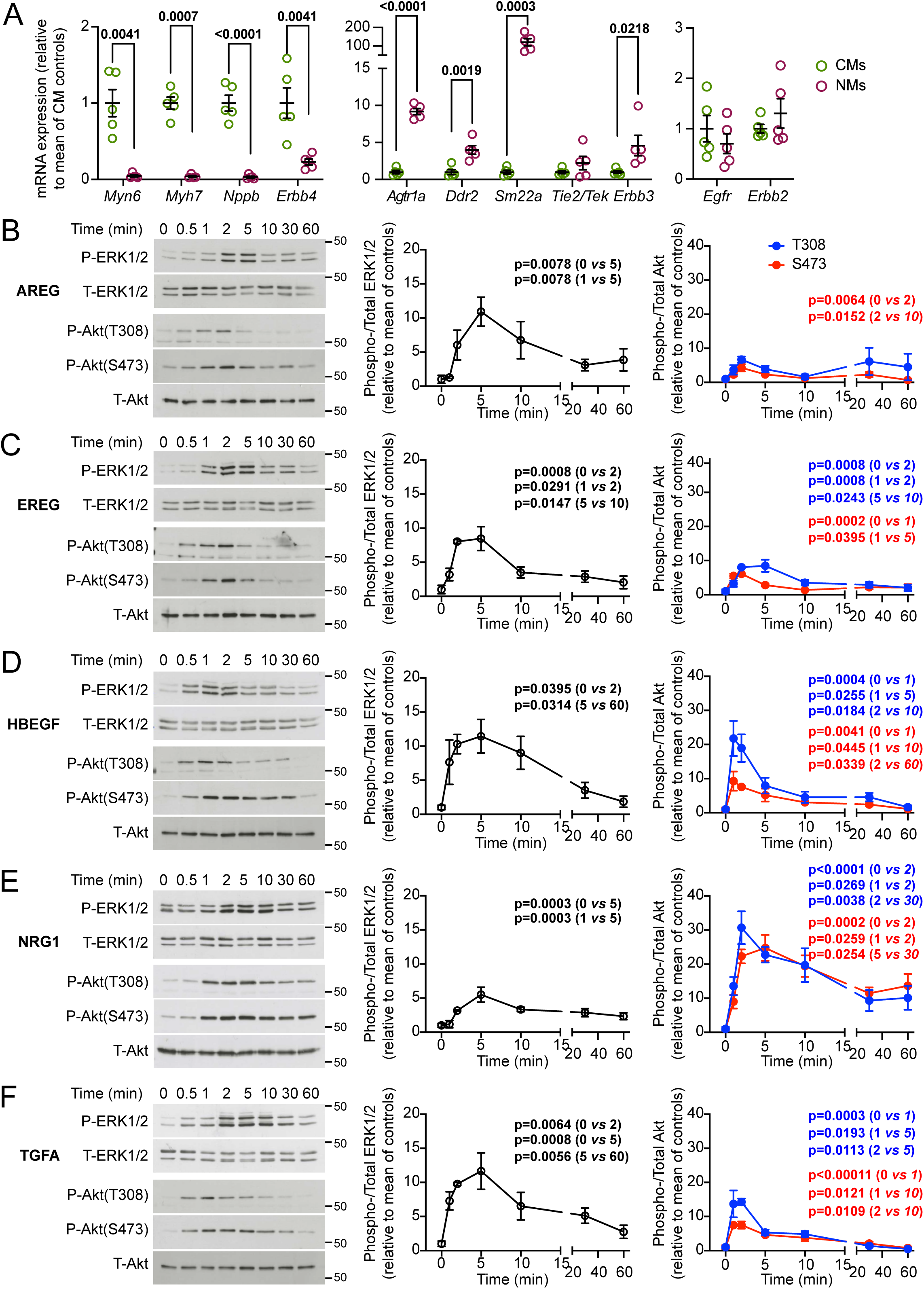
EGFR family ligands promote phosphorylation of ERK1/2 and Akt to different degrees in rat neonatal cardiomyocytes. **A**, mRNA expression of EGFR family members in neonatal rat cardiomyocytes (CMs) and cardiac non-myocytes (NMs) compared with markers for cardiomyocytes (left panel) and cardiac non-myocytes (centre panel). **B-F**, Phosphorylation (i.e. activation) of ERK1/2 and Akt in rat neonatal cardiomyocytes. Cells were treated with 50 ng/ml AREG (**A**), EREG (**B**), HBEGF (**C**), NRG1 (**D**) or TGFA (**E**) for the indicated times. Proteins were immunoblotted with antibodies to phosphorylated (P) or total (T) ERK1/2 or Akt. Representative blots are on the left with densitometric analysis for ERK1/2 and Akt in the middle and right panels, respectively. Densitometric data are means ± SEM (n=4 independent cardiomyocyte preparations) and are the ratio of phosphorylated:total protein, normalised to the mean of the zero-time samples. Over 80% of treatments (each time point for each agonist) met conditions of normality (Shapiro-Wilk tests). Statistical analysis used one-way ANOVA with Tukey’s multiple comparisons post-hoc tests. Selected p values are shown for P/T ERK1/2 (black type), P/T Akt(T308) (blue type) and P/T Akt(S473) (red type) to illustrate significant increases and decreases in phosphorylation.

EGFR-Ls elicit different phenotypic effects [22] which may reflect differences in duration and intensity of the intracellular signalling. EGF (a high affinity ligand for EGFRs) activates ERK1/2 and Akt in cardiomyocytes [30]. Here, we compared the profiles of ERK1/2 and Akt phosphorylation in rat neonatal cardiomyocytes treated with TGFA and AREG (high and low affinity ligands for Egfrs, respectively), EREG and HBEGF (low and high affinity ligands for Egfrs and ERBB3, respectively) and NRG1 (that activates ERBB4 and ERBB3), using antibodies to the activating phosphorylations for immunoblotting. All ligands activated both signalling pathways with maximal activation within 2-5 min, and the response was transient such that much of the response was lost by 60 min (**Figure 2B-F**). At 50 ng/ml (a concentration that gives maximal or close to maximal activation; **Supplementary Figure S1**), the degree of activation of ERK1/2 by AREG, EREG, HBEGF and TGFA was similar and appeared higher than that induced by NRG1. In contrast, NRG1 potently activated Akt with lesser activation by the other ligands. Thus, activation of Erbb4 in cardiomyocytes had a dominant effect on Akt signalling, whilst activation of EGFR had a more pronounced effect on ERK1/2. Close inspection of the profiles identified differences between high and low affinity ligands for Egfrs. High affinity ligands (HBEGF, TGFA) promoted very rapid phosphorylation of ERK1/2 and Akt within 30 s that was maximal at ∼2 min, whereas low affinity ligands (AREG, EREG) promoted slower phosphorylation of ERK1/2 and limited activation of of Akt.

EGFR and ERBB4 can operate as heterodimers with ERBB2. To determine the degree to which EGFR and ERBB4 signalling is mediated via ERBB2, cardiomyocytes were treated with EGF (to activate Egfrs) or NRG1 (to activate Erbb4) (100 ng/ml, 5 min) without/with small molecule inhibitors. Afatinib was used at 1 µM to inhibit all EGFR tyrosine kinases (IC_50_ values for ERBB2 and EGFR are 0.5 nM and 14-16 nM, respectively) [52, 53] and 1 µM tucatinib was used for selective inhibition of ERBB2 [60]. We compared the response to ET-1 which induces maximal activation of ERK1/2 but minimal activation of Akt [27, 28, 30]. EGF and NRG1 activated ERK1/2 to a similar degree, but less than ET-1 (**Figure 3A-B**). As expected, afatinib attenuated activation by EGF or NRG, but had no significant effect on the activation of ERK1/2 by endothelin-1. Tucatinib had no significant effect on activation of ERK1/2 by EGF, but partially inhibited the response to NRG1 (50.4% inhibition). NRG1 activation of Akt was greater than that induced by EGF (26.7 ± 3.2 fold *vs* 7.6 ± 2.3 fold relative to vehicle only; means ± SEM, n=4), whilst ET-1 did not promote any significant phosphorylation of Akt (**Figure 3B**). Activation of Akt by EGF or NRG1 was inhibited by afatinib, whilst tucatinib inhibited activation of Akt by NRG1 (73.4% inhibition). Thus, EGF has no significant requirement for ERBB2 to activate ERK1/2 or Akt, whereas more of the NRG1 signal is via ERBB2.

**Figure 3.**
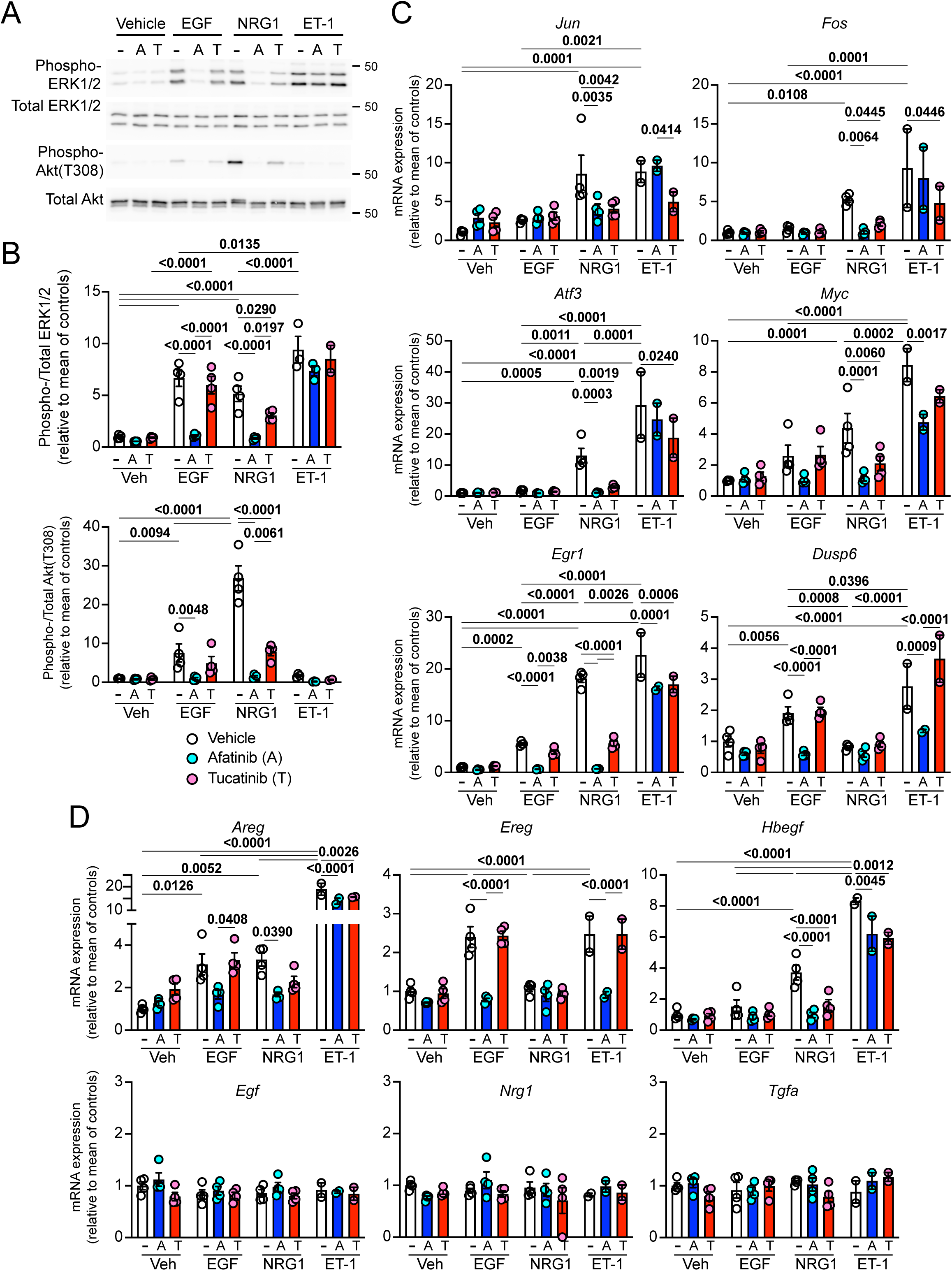
Effects of afatinib *vs* tucatinib on ERK1/2 and Akt phosphorylation and immediate early gene expression induced by EGF and NRG1 in cardiomyocytes. Rat neonatal cardiomyocytes were treated with EGF or NRG1 (100 ng/ml, 5 min) in the absence or presence of 1 µM afatinib or tucatinib. **A-B**, Proteins (10^5^ cells per sample) were immunoblotted for phosphorylated (phospho-) or total ERK1/2 or Akt. **A**, Representative blots. **B**, Densitometric analysis. **C-D**, RNA was extracted and mRNA expression of selected immediate early genes (**C**) or EGFR family ligands (**D**) was measured by qPCR. **B-D**, Data are means ± SEM (n=4 independent cardiomyocyte preparations; individual data points shown) and are normalised to the mean of vehicle treated controls. Statistical analysis used two-way ANOVA with Tukey’s multiple comparisons post-hoc tests. p values for significant changes (p<0.05) are in bold type.

ERK1/2 and Akt promote phosphorylation and activation of different transcription factors to modulate gene expression [32]. It is therefore likely that activation of EGFRs by EGF elicits different changes in gene expression compared with activation of ERBB4 by NRG1. We next used qPCR to determine the downstream consequences of EGF and NRG1 signalling on gene expression, focusing on upregulation of immediate early genes at 1 h [27]. NRG1, but not EGF, promoted significant increases in expression of *Jun*, *Fos*, *Atf3* and *Myc* mRNAs and this upregulation was abolished by afatinib or tucatinib (**Figure 3C**). The increase in expression of *Egr1* induced by NRG1 was greater than that induced by EGF; afatinib attenuated this increase by either agonist but tucatinib only inhibited the response to NRG1. mRNA expression of *Dusp6* (an inducible dual specificity phosphatase that is upregulated by and dephosphorylates/inactivates ERK1/2 [61]) was increased by EGF but not NRG1, a response that was inhibited by afatinib, but not tucatinib. We compared the effects of the two ligands on expression of EGFR-Ls (**Figure 3D**). Whereas *Areg* was similarly upregulated by both agonists, EGF (not NRG1) increased expression of *Ereg* whilst NRG1 (not EGF) increased expression of *Hbegf*. Increased expression of *Areg* or *Hbegf* by NRG1 were both inhibited by tucatinib. For all of these genes, expression was increased by ET-1 to a similar or greater extent as EGF or NRG1. Despite the lack of effect on ERK1/2 or Akt phosphorylation (**Figure 3A-B**), the increase in expression of all genes except *Ereg* was reduced to some degree when cells were co-treated with afatinib or tucatinib (**Figure 3C-D**). Thus, it is possible that ET-1 signalling to gene expression has some requirement for EGFRs. *Egf*, *Nrg1* and *Tgfa* mRNA expression was unaffected by any of the treatments. These data highlight the mechanistic complexities relating to the regulation of mRNA expression, demonstrating differential effects of EGF and NRG1, and indicating that ERBB2 plays a prominant role in the response to NRG1.

To further examine the differences between responses to EGF and NRG1, selected samples were analysed using RNASeq (vehicle, tucatinib, EGF, NRG1 and EGF or NRG1 with tucatinib; n=4 per group). Similar numbers of differentially expressed genes (DEGs) were identified as upregulated by EGF and NRG1 with fewer DEGs downregulated by EGF than NRG1; 223 and 136 transcripts were upregulated and downregulated, respectively, by either EGF or NRG1 (**Figure 4A, Supplementary Figure S2, Supplementary spreadsheet SS1**). Tucatinib alone promoted changes in expression of 125 DEGs, suggesting that ERBB2 maintains expression of some genes in the basal state. Tucatinib significantly inhibited the changes of each of 100 of the 815 DEGs that were up- or downregulated by EGF (12.3%) and 455 of 995 DEGS up- or downregulated by NRG1 (45.7%). In addition, for genes not significantly inhibited by tucatinib on an individual basis, as a group, there was still a small but significant inhibition of mRNAs that were downregulated by either EGF or NRG1 and for those which were upregulated by NRG1. qPCR results for the selected immediate early genes and EGFR-Ls studied in **Figure 4C-D** were confirmed in the RNASeq dataset (**Supplementary Figure S3**).

**Figure 4.**
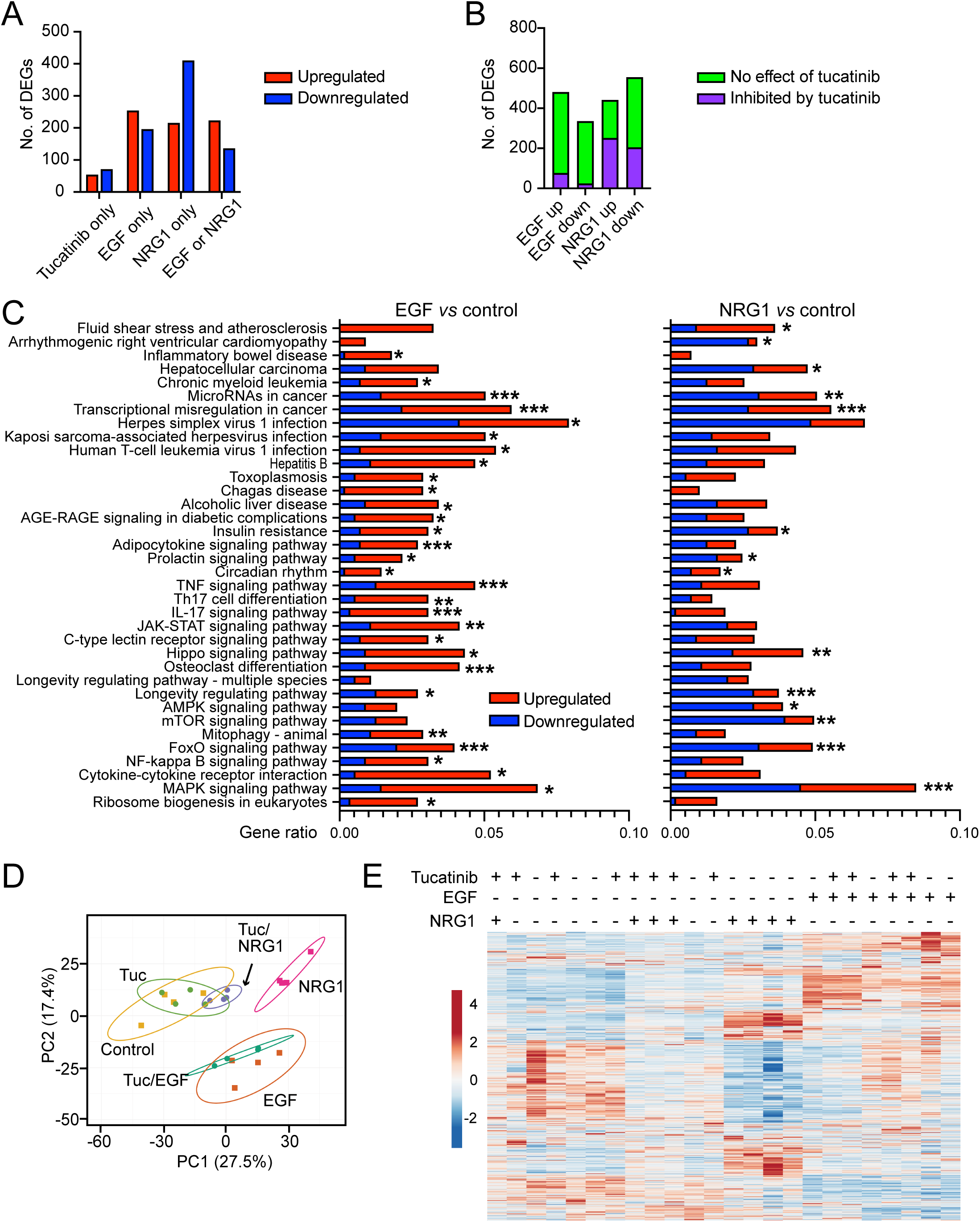
Inhibition of ERBB2 with tucatinib differentially affects the changes in the cardiomyocyte transcriptome induced by EGF *vs* NRG1. Rat neonatal cardiomyocytes were treated with EGF or NRG1 (100 ng/ml, 60 min) in the absence or presence of 1 µM tucatinib and mRNA expression measured by RNASeq (n=4 per group). Differentially expressed genes (DEGs) were identified (FDR<0.05). **A**, Numbers of upregulated (red) and downregulated (blue) DEGs identified with tucatinib, EGF or NRG1 relative to control cells. **B**, Genes upregulated (up) or downregulated (down) by EGF or NRGs unaffected by or significantly inhibited by tucatinib were identified from the RNASeq dataset. **C**, KEGG enrichment analysis for upregulated (red) and downregulated (blue) genes for significantly changed categories identified with EGF (left) or NRG1 (right) demonstrates qualitatively different responses induced by the two ligands. * p<0.05, * p<0.01, *** p<0.005 (adjusted p value; see Methods section for details and **Supplementary Spreadsheet SS3** for gene identities and individual p values). **D**, PCA analysis of treatment conditions by ClustVis. Unit variance scaling is applied to rows; SVD with imputation is used to calculate principal components. X and Y axes show principal components 1 and 2 that explain 27.5% and 17.4% of the total variance, respectively. Prediction ellipses are such that with probability 0.95, a new observation from the same group will fall inside the ellipse. **E**, Heatmap for differentially expressed genes constructed using ClustVis. Rows are centered; unit variance scaling is applied to rows. Both rows and columns are clustered using correlation distance and complete linkage.

To determine if the overall changes in gene expression induced by EGF *vs* NRG1 were qualitatively different, we used KEGG analysis of the DEGs identified with EGF compared with NRG1. This illustrated different patterns of gene regulation from the two agonists which is likely to result in different phenotypic consequences (**Figure 4C; Supplementary spreadsheet SS3**). For example, NRG1 had a greater effect on genes associated with AMPK or mTOR signalling (pathways linked to metabolism, protection and aging [62, 63]) than EGF, whereas EGF had a greater effect on immune cell signalling pathways (TNF, IL17, JAK-STAT signalling). Even for pathways that were significantly affected by both agonists (e.g. MAPK signalling) the responses differed in the proportions of upregulated *vs* downregulated genes. PCA analysis and hierarchical clustering showed that cells treated with tucatinib alone or tucatinib with NRG1 were most similar to vehicle only controls and cells treated only with NRG-1 clustered away from these samples (**Figure 4D-E**). This indicates that the global response to NRG1 was substantially affected by tucatinib, indicating that they result from ERBB2 activation. As expected, cardiomyocytes treated with NRG1 *vs* EGF clustered apart. However, samples treated with EGF with or without tucatinib clustered more closely together indicating that there was much less effect of tucatinib on the overall EGF response. In conclusion, the RNASeq data validate on a global level the key conclusions from the qPCR data that more of the effects of NRG1 on cardiomyocyte gene expression require ERBB2 than EGF, but also demonstrate that the downstream effects of EGF on gene expression are qualitatively different from those induced by NRG1.

### Afatinib inhibits cardiac hypertrophy induced by AngII and enhances cardiac fibrosis

The acute changes in mRNA expression of EGFR-Ls in cardiomyocytes induced by hypertrophic stimuli (**Figure 3D**; [27, 28]) suggest that the network is likely to be involved in cardiac hypertrophy *in vivo*. To assess this, we used a model of cardiac hypertrophy induced by a “slow-pressor” dose of angiotensin II (AngII, 0.8 mg/kg/d) in young male C57Bl/6J mice to induce an adaptive response over 3-7 d without substantial changes in ejection fraction or cardiac output (as in [64, 65]). *Areg*, *Ereg*, *Hbegf* and *Tgfa* mRNAs were upregulated in mouse hearts after 3 d treatment with AngII, whilst *Egf* mRNA was downregulated (**Figure 5A**), supporting the concept that EGFR-Ls are important as cardiac hypertrophy develops. To determine if this is the case, we assessed the effects of afatinib (at a concentration to inhibit all EGFRs in the 3-7 d period of the study; see Methods for details) on cardiac hypertrophy induced by AngII over 7 d (**Figure 5B**). Mice were treated with vehicle, 0.8 mg/kg/d AngII or AngII with 4.5 mg/kg/d afatinib. Cardiac function/dimensions were monitored by echocardiography with analysis of long axis B-mode images of the heart using speckle-tracking (strain analysis) (**Figure 5C-D; Supplementary Table S6**). Over 7 d, AngII alone had no significant effect on established measures of global cardiac function (fractional shortening, ejection fraction, stroke volume and cardiac output) or on global longitudinal strain (GLS, an early marker of cardiac dysfunction [66]) (**Figure 5E**), consistent with our recent studies [65]. AngII did not significantly affect LV end diastolic volume, but increased predicted LV end diastolic mass (LVEDM) consistent with adaptive cardiac hypertrophy. Inclusion of afatinib with AngII reduced the increase in LVEDM compared with AngII alone, with a significant decline in GLS at 3 d, and signs of decreasing stroke volume and cardiac output at 7 d.

**Figure 5.**
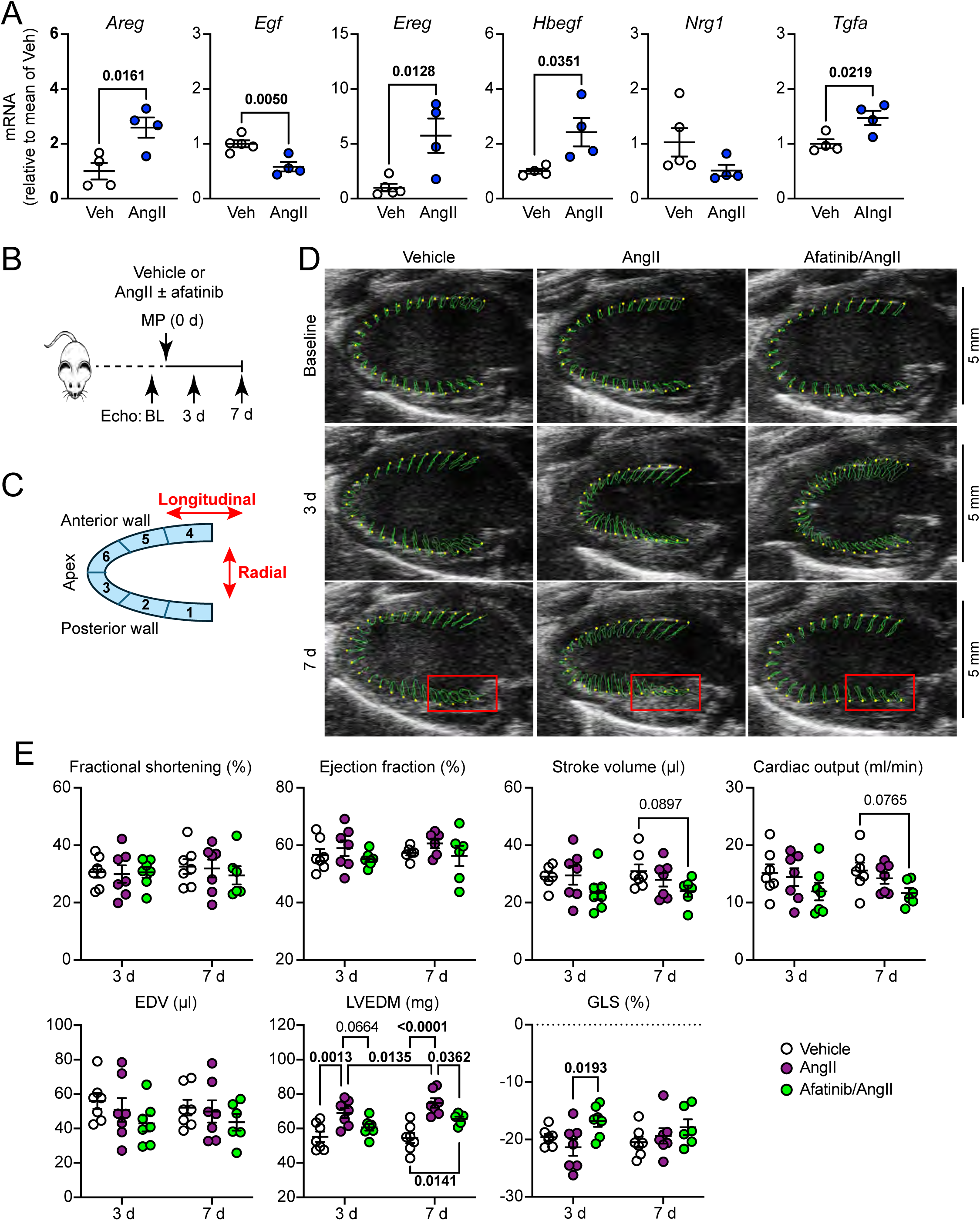
Afatinib reduces cardiac hypertrophy induced by AngII. **A**, mRNA expression of EGFR family ligands in hearts from male C57Bl/6J mice treated with vehicle (Veh) or 0.8 mg/kg/d AngII (3 d). Data are means ± SEM (n=4 or 5) with individual data points shown. Statistical analysis used unpaired t tests. **B**, Schematic of experiment with AngII ± afatinib. Male C57Bl/6J mice (9 weeks) received a baseline (BL) echocardiogram (Echo) with subsequent echocardiograms at 3 and 7 d after implantation of osmotic minipumps (MP) to deliver vehicle (Veh), 0.8 mg/kg/d AngII or AngII with 4.5 mg/kg/d afatinib (n=7 per group). **C**, Diagram of left ventricle showing longitudinal and radial vectors, with identification of segments for analysis. **D**, Representative long axis B-mode images for a single heart for each condition. The green line traces the movement of the endocardial wall (identified by the yellow spots) across two cardiac cycles. **E**, Speckle-tracking (strain) analysis of long axis B-mode images showing global measures of cardiac function (fractional shortening, ejection fraction, stroke volume, cardiac output) and dimensions (EDV, end diastolic volume; LVEDM, left ventricular end diastolic mass; GLS, global longitudinal strain). Individual values are shown with means ± SEM. Statistical analysis used two-way paired (according to time) ANOVA with Tukey’s multiple comparisons post-hoc tests. p values for significant changes (p<0.05) are in bold type; values of 0.05-0.1 in standard type.

Speckle-tracking software traces the endocardial wall and tracks movement in longitudinal and radial directions, in line with and perpendicular to the endocardial wall, respectively, and can be used to assess movement of different segments around the LV (**Figure 5C**). Segmental analysis of LV endocardial movement identified variation around the LV, with particularly limited longitudinal movement at the apex (segments 3 and 6) compared with the mid- and basal regions (segments 2 and 5, and 1 and 4, respectively) (**Figure 6A, Supplementary Spreadsheet SS2**) (as in [65]). Inclusion of afatinib with AngII caused significant reductions in peak longitudinal displacement and strain in segments 1 and 4 in the basal region of the heart compared with AngII alone (3 d or 7 d) with some effect in segments 2 and 5 (mid-regions) at 3 d (**Figure 6B-C**). The effects on segments 1 and 4 were more pronounced than global strain, GLS (**Figure 5E**), presumably because GLS includes data from the relatively unaffected apical region. Examination of segment 1 on the echocardiograms particularly illustrated the effects on radial *vs* longitudinal movement of AngII vs afatinib/AngII (**Figure 5C**, red boxes).

**Figure 6.**
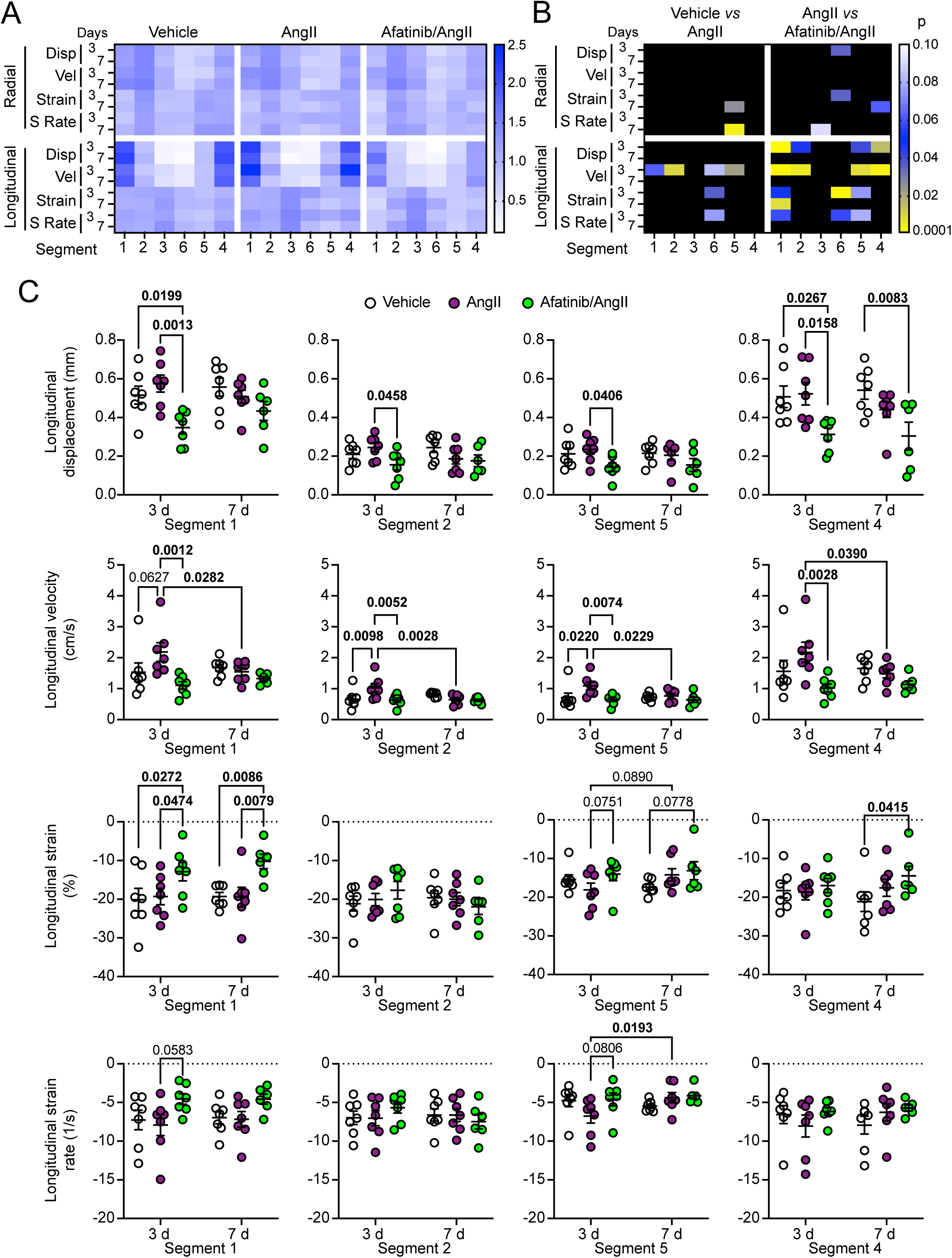
Afatinib promotes cardiac dysfunction in selected segments of the left ventricular endocardium in mice treated with AngII. **A**, Heatmaps showing mean values for each parameter assessed at 3 or 7 d for each of the 6 segments shown in Figure 5C. Data were normalised to the mean for all the segments for each parameter. Parameters assessed were peak values for radial and longitudinal movement, assessing displacement (Disp), velocity (Vel), strain and strain rate (S Rate). **B**, Heatmaps showing statistical probabilities for differences in each parameter between mice treated with vehicle *vs* AngII (left) or AngII *vs* AngII with afatinib (right). A two step gradient was used: 0.001-0.05, yellow-blue; 0.05-0.10, blue-white. **C**, Peak longitudinal displacement, velocity, strain and strain rate at 3 d and 7 d. Individual values are shown with means ± SEM. **B-C**, Statistical analysis used two-way paired (according to time) ANOVA with Tukey’s multiple comparisons post-hoc tests. p values for significant changes (p<0.05) are in bold type; values of 0.05-0.1 in standard type.

Histological analysis of hearts collected from a subgroup of mice at 7 d demonstrated that AngII increased cardiomyocyte cross sectional area (i.e. hypertrophy) and this was reduced by addition of afatinib (**Figure 7A**). However, expression of *Myh7* (a marker of hypertrophy [15]) was enhanced in hearts from mice treated with afatinib and AngII (**Figure 7B**), presumably a reflection of ongoing or enhanced stress. Cardiac fibrosis was increased by AngII, particularly in the perivascular region and this was increased further in hearts from mice treated with afatinib and AngII (**Figure 7C**). AngII also increased expression of mRNAs for collagens, other extracellular matrix proteins (*Fn1*, *Postn*), and other pro-fibrotic proteins [*Ccn2* (connective tissue growth factor, CTGF), *Tgfb1*], but this was unaffected by addition of afatinib (**Figure 7D, Supplementary Figure S4**), suggesting that the pro-fibrotic effects of afatinib were not directly linked to increased transcription. We assessed the effects of afatinib on expression of mRNAs for EGFR-Ls. The changes in expression of *Areg*, *Egf* and *Tgfa* detected at 3 d in hearts from mice treated with AngII (**Figure 5A**) were not detected at 7 d, whilst the small increases in expression of *Ereg* and *Hbegf* were essentially unchanged by addition of afatinib (**Figure 7E**). Interestingly, whilst AngII alone did not significantly affect expression of *Nrg1* mRNA, *Nrg1* was significantly reduced by treatment with afatinib and AngII. Neither AngII nor afatinib plus AngII significantly affected expression of mRNAs for the EGFRs (**Figure 7F**).

**Figure 7.**
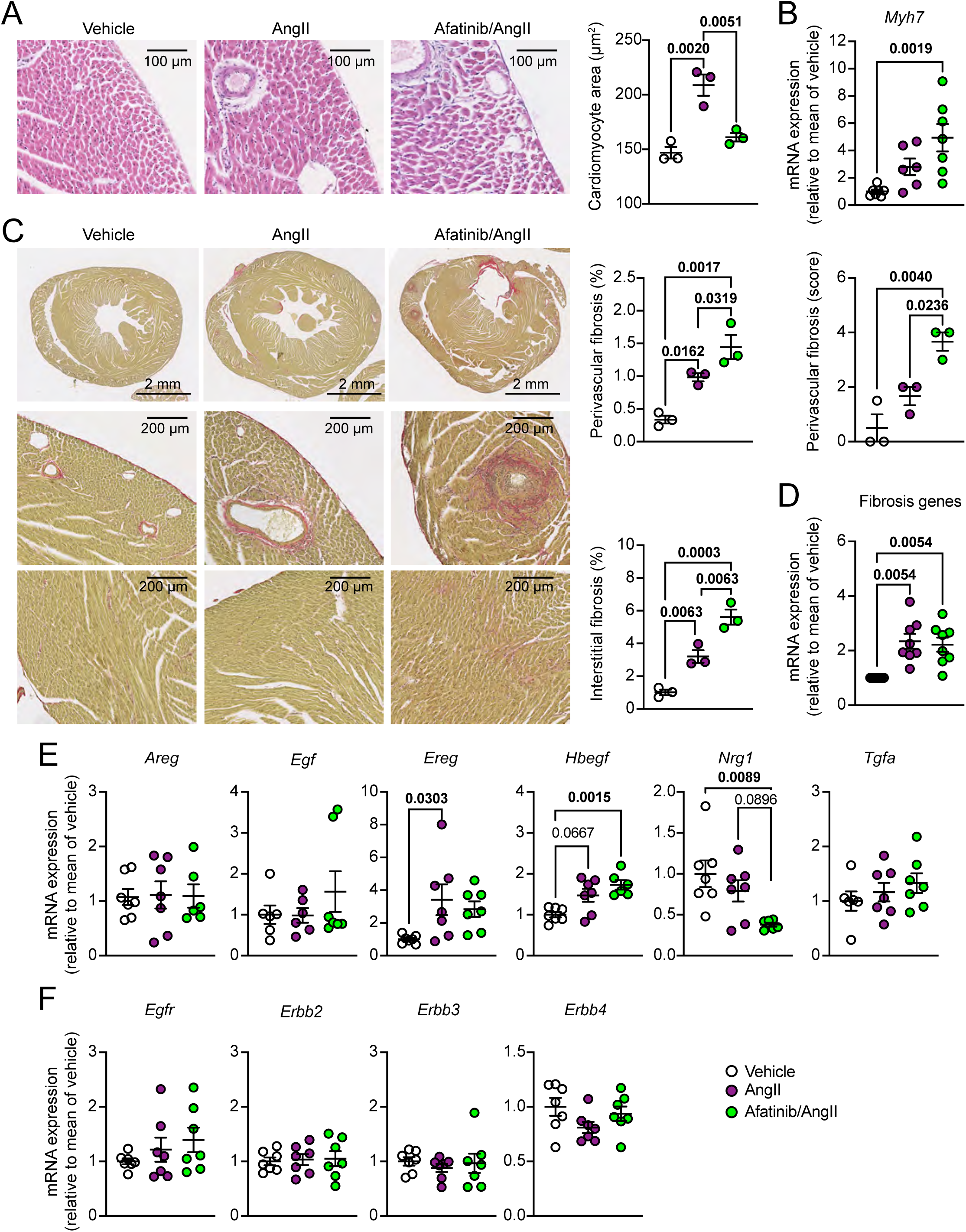
Afatinib inhibits cardiomyocyte hypertrophy induced by AngII in mouse hearts but enhances cardiac fibrosis. Male C57Bl/6J mice were treated with vehicle, 0.8 mg/kg/d AngII or AngII with 4.5 mg/kg/d afatinib for 7 d. Mice were culled, and hearts were fixed in formalin for histological analysis (**A** and **C**) or ground to powder under liquid N_2_ and samples used to prepare RNA for qPCR (**B, D-F**). **A**, H&E staining was used to assess cardiomyocyte area. Representative images are on the left with analysis shown on the right. **B**, Picrosirius red staining to show fibrosis. Representative images are on the left with analysis shown on the right. **B, D-E**, qPCR analysis for *Myh7*, a marker of hypertrophy (**B**), the mean of selected fibrosis genes (*Col1a1*, *Col2a1*, *Col3a1*, *Col4a1*, *Fn1*, *Postn1*, *Cnn2*, *Tgfb1*) (**D**), EGFR family ligands (**E**) and EGFR family members (**F**). Statistical analysis used one-way ANOVA with Tukey’s post-hoc multiple comparisons tests. p values for significant changes (p<0.05) are in bold type; values of 0.05-0.1 in standard type.

These data suggest the EGFR network plays an important role in cardiac hypertrophy induced by AngII. Not only did AngII stimulate mRNA expression of EGFR-Ls from 3 d, but subsequent cardiac hypertrophy induced by AngII was moderated by afatinib with reduced cardiomyocyte hypertrophy and enhanced cardiac fibrosis, and effects on cardiac function detected with afatinib and AngII compared with AngII alone.

## Discussion

“Mixed signals in heart failure: cancer rules”, the title of a 2002 review by Hoshijima and Chien [67] is as pertinent now as it was then. Many of the key intracellular signalling pathways that drive cancer, also maintain cardiac homeostasis or drive adaptation and disease. As in cancer, development of cardiac diseases results from intercellular mediators activating protein kinase signalling leading to phosphorylation of transcriptional regulators and changes in gene expression. Similarities between the diseases extend beyond this acute phase which lasts just minutes to hours with, for example, initial tumour formation or cardiac hypertrophy developing over days to weeks, and phenotypic end-points (metastatic disease or heart failure) coming after months or years of continued stress and signalling. Like cancer, heart failure is a multi-hit disease with potential for prevention, even reversal, at any step along the way. However, the step markers for cancer are more clearly defined than those of heart failure, and deeper understanding of the molecular mechanisms is required for the heart. With a focus only on the EGFR system, we highlight many of the principles raised by Hoshijima and Chien (**Figure 8**) with acute activation of intracellular signalling in cardiomyocytes (seconds to minutes; **Figures 2** and **3**), modulation of gene expression (minutes to hours; **Figures 3** and **4**), upregulation of EGFR-Ls in the early phase of developing cardiac hypertrophy induced by AngII in mice with modulation of the hypertrophic response by an inhibitor of the EGFR system (days to weeks; **Figures 5** and **7**), and changes in expression of EGFR-Ls in human heart failure (months to years; **Figure 1**). Thus, the EGFR system is implicated at every step along the way to heart failure.

**Figure 8.**
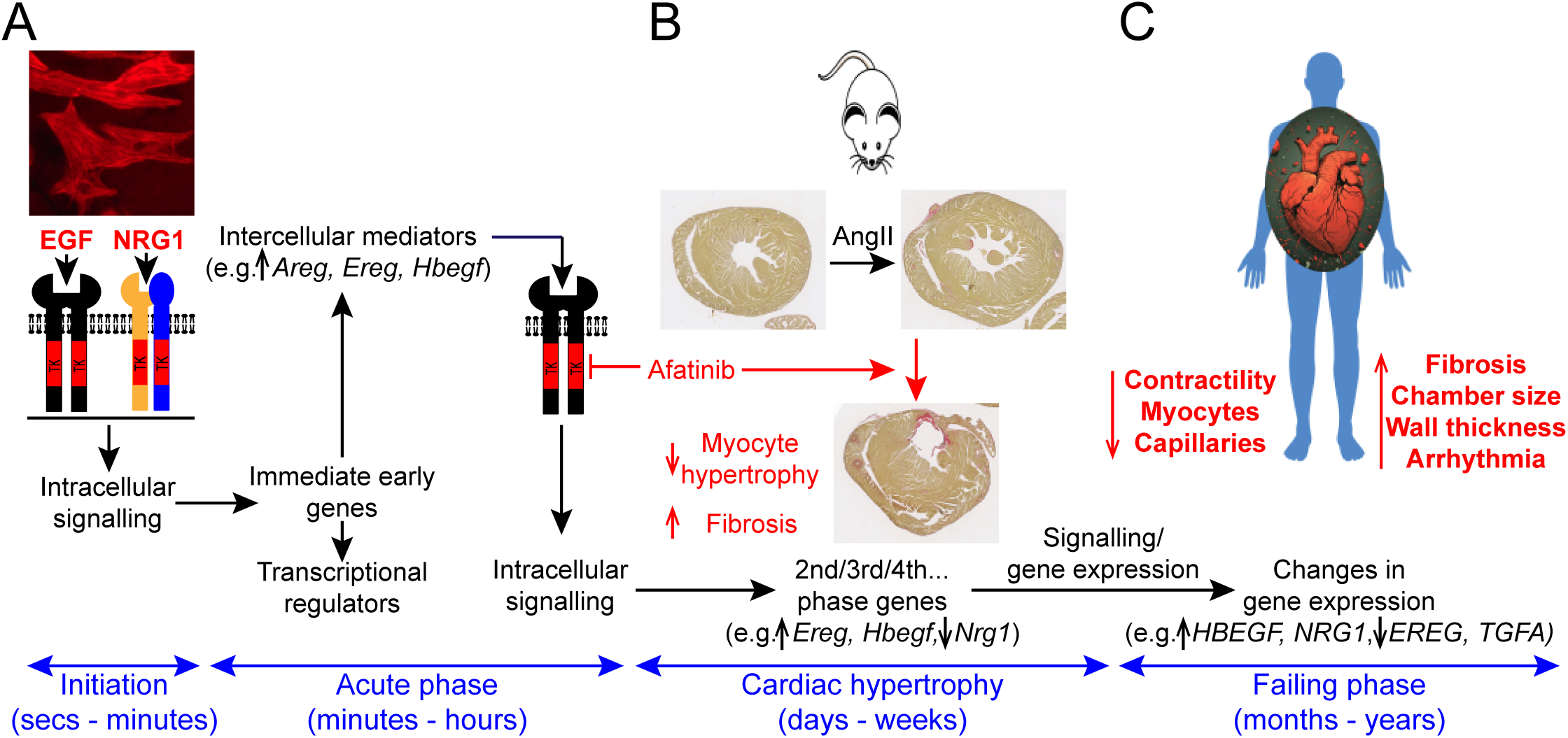
Principles of signalling to gene expression and cardiac disorder applied to the EGFR network. Development of cardiac diseases initiates from pathophysiological stresses leading to production of intercellular signals such as EGFR family ligands (EGFR-Ls). **A**, EGFR-Ls binding to cell surface receptors on cardiomyocytes rapidly initiated intracellular signalling (seconds to minutes) leading to changes in gene expression over minutes to hours (acute phase). Changes in expression of transcriptional regulators are predicted to modulate future transcriptional responses whilst production of additional growth factors (e.g. *Areg*, *Ereg* and *Hbegf*) can propagate the intercellular signal. **B**, Continued stress and multiple rounds of signalling/gene expression over days to weeks result in adaptive cardiac hypertrophy, as demonstrated in a mouse model of cardiac hypertrophy induced by angiotensin II (AngII). Inhibiting EGFR family signalling with afatinib at this stage reduced the degree of cardiomyocyte hypertrophy but increased cardiac fibrosis, highlighting the role of the EGFR network in disease development. This was associated with further changes in gene expression including some EGFR-L mRNAs. **C**, Over many months and years, continued pathophysiological stresses with ever changing patterns of signalling and gene expression result in heart failure with features such as decreased contractility, loss of cardiomyocytes, a reduced capillary network and increased fibrosis. Changes in expression of EGFR-Ls were detected in human failing hearts, further implicating the EGFR network in end-stage disease. Thus, the EGFR system is implicated at every step along the way to heart failure. Intervention in this system offers opportunities to alleviate disease progression and/or facilitate rescue.

### The cardiac EGFR network and intercellular cardiac signalling

Although development of cardiac disease and cancer are associated with many of the same signalling pathways, the cells are fundamentally different. Cancer cells have high plasticity resulting from disease-causing mutations and ongoing mutational adaption. In contrast, terminally-differentiated cardiomyocytes are “hard-wired” with respect to intracellular signalling; mutations do not accumulate and signalling pathways remain robust. The heart contains other cells with proliferative potential (e.g. endothelial cells, smooth muscle cells, cardiac fibroblasts). Since cardiac tumours are rare [68], it may be assumed cell division and signalling networks are highly constrained even in these cells. The signalling pathways (like the ERK1/2 cascade and Akt) are similar, but inputs and outputs differ.

Inputs from intercellular signals (e.g. EGFR-Ls) trigger cell surface receptors (e.g. EGFR family members). Profiling receptor expression across cardiac cell types is a first stage in predicting which cells will respond to different stimuli. The dogma for EGFR family members in the heart is that cardiac endothelial cells and fibroblasts express all EGFR family members, whereas cardiomyocytes express only ERBB2 and ERBB4 [39]. Single cell sequencing data from the HPA did, indeed, indicate greater expression of *ERBB2* and *ERBB4* than other receptors in cardiomyocytes with almost exclusive expression of *ERBB4* in these cells, but expression levels of *ERBB4* were ∼10-fold greater than those of *ERBB2* (**Figure 1B**, upper panel). Expression of *EGFR* was predominantly in cardiac non-myocytes, whilst expression of *ERBB2* and *ERBB4* varied according to cell type. It is more difficult technically to assess protein expression in individual cells than mRNA, particularly for cell surface transmembrane receptors. However, a particularly informative proteomics study focused on cell surface glycoproteins in cardiac cells (surfaceome mapping) [22], identifying expression of ERBB2/ERBB4 in cardiomyocytes, EGFR/ERBB2 in cardiac fibroblasts, ERBB2 in endothelial cells and EGFR/ERBB2 in smooth muscle cells. Allowing for lower sensitivity of proteomics, the data indicate that, at least for the EGFR family members, transcriptomics data are reasonably predictive of protein expression. However, proteomics data have yet to shed light on the ratios of different receptors in individual cells as indicated from transcriptomics studies.

Intercellular signals are as important in maintaining homeostasis as in driving adaptation, so profiling ligand expression across cell types is informative. HPA data suggest that autocrine and paracrine systems operate for the EGFR network in the heart (**Figure 1B**, lower panel) with, perhaps, paracrine signalling of EGF from cardiomyocytes to EGFR receptors in cardiac non-myocytes and NRG1 from endothelial cells to ERBB4 in cardiomyocytes. Identification of other neuregulins (*NRG2*, *NRG3* and *NRG4*) in cardiac cells, particularly cardiomyocytes, suggests they also play an important autocrine/paracrine role. Less is known of the roles of these neuregulins in the heart. NRG4 has been best studied as an adipokine released into the circulation from brown adipose tissue with cardioprotective properties [39], but the roles of NRG2 and NRG3 in the heart remain to be established. Local endogenous expression of peptide growth factors for the EGFR family is difficult to study at the protein level using conventional methods, because of their size and low overall levels of expression (though high local levels can accumulate). Studies in other cells have used molecular engineering to generate fluorescent forms of EGFR-Ls to study their effects on epithelial cell migration [69]. Such methodologies remain to be applied to the cardiac system.

Changes in ligand and/or receptor profiles allows cells to detect and respond to a changing environment (e.g. an increase in workload for the heart). We did not detect significant changes in expression of mRNAs for the EGFRs in any of the systems we studied (**Figure 1C**), but other studies indicate that ERBB2 protein is downregulated in human failing hearts, and ERBB4 may be up- or downregulated [18, 19, 22]. The difference may reflect post-transcriptional/translational regulation of receptor expression including increased turnover but, equally, there may be differences in patient populations or aetiology of disease. Although we did not identify changes in receptor expression, there were significant changes in mRNAs for EGFR-Ls in neonatal cardiomyocytes treated with EGF, NRG1 or ET-1 (upregulation or *Areg*, *Ereg* and/or *Hbegf*), in hearts from mice treated with AngII (upregulation of *Areg*, *Ereg*, *Hbegf* and *Tgfa* at 3 d; upregulation of *Ereg* and *Hbegf* at 7 d) and in human failing hearts (upregulation of *HBEGF*, *NRG1* and possibly *EGF*; downregulation of *EREG* and *TGFA*). Upregulation of *Hbegf* in all three systems suggests it plays an important role in cardiac hypertrophy and heart failure. This is consistent with other studies in which *HBEGF* mRNA is upregulated in hypoxic vs normoxic biopsies from human hearts [70]. Studies of overexpressed Hbegf in rat neonatal cardiomyocytes and mouse hearts support the concept that it operates in a local environment to promote cardiomyocyte hypertrophy via autocrine/paracrine signalling [71], so increases in mRNA expression are likely to be relevant. However, HBEGF itself is also increased in plasma from patients with cardiac hypertrophy or atrial fibrillation [72, 73] which further highlights its potential importance. For other EGFR-Ls, there was less consistency, probably because of temporal differences in gene regulation. The data for *EREG/Ereg* are particularly interesting because it was downregulated in end-stage disease but upregulated in cardiomyocyte treated with EGF (not NRG1) and in hearts of mice treated with AngII at 3 or 7 d. This profile suggests it may be needed to drive/support adaptation, in which case downregulation in end-stage disease may be a detrimental factor. Consistent with this, *Ereg* is upregulated in a rat model of acute myocardial infarction (7 d) and downregulation of *Ereg* impairs angiogenesis causing enhanced cardiac dysfunction [74]. Further studies of EGFR-Ls in the heart will be important to dissect the phenotypic effects of the network.

### The cardiac EGFR network and intracellular signalling to gene expression

ERBB4 receptors are activated by 4 different neuregulins, whilst EGFRs are activated by 7 different ligands (**Figure 1A**) of high or low affinity (EGF, TGFA and HBEGF being high affinity ligands). Activation of ERK1/2 and Akt by these ligands varied substantially. Neuregulin binding to ERBB4 in the heart is recognised to activate Akt [39] but, although NRG1 potently activated Akt in neonatal cardiomyocytes, it also activated ERK1/2 (**Figures 2** and **3**). In contrast, ligands for the EGFR preferentially activated ERK1/2 rather than Akt. Different ligands binding to the EGFR have different phenotypic consequences [22], and the purpose of the experiments in **Figure 2** and **Supplementary Figure S1** was to determine if this was reflected at the level of ERK1/2 or Akt. Indeed, the profiles for activation of ERK1/2 and Akt by high affinity ligands (HBEGF, TGFA) differed from lower affinity ligands (AREG, EREG) with more rapid phosphorylation of ERK1/2 and Akt by the former and slower kinetics for ERK1/2 phosphorylation with limited Akt phosphorylation for the latter. Much (but not all) of the signal was lost within 60 min, indicating that the response was transient. However, further information relating to the location of the signal may be important. For example, if residual activity at 60 min is retained in the nucleus, this could be sufficient for continued effects on gene expression. The biological significance of variations in signal intensity and duration remains to be fully investigated in cardiomyocytes but the most immediate effect is likely to be on immediate early gene expression.

ERK1/2 and Akt signalling promotes phosphorylation of transcriptional regulators which triggers an immediate early gene response as shown here in cardiomyocytes (**Figures 3** and **4**) and in our previous studies of hypertrophic agonists in cardiomyocytes and perfused hearts [27, 28, 43]. Since ERK1/2 and Akt activate different transcription factors [32], the profile of immediate early genes regulated by each cascade will differ. Consequently, the differences in gene expression induced by EGF vs NRG1 in cardiomyocytes were not unexpected given the preferences of these ligands for activation of ERK1/2 *vs* Akt, respectively (**Figures 2, 5** and **6**; **Supplementary Figure S2**). However, the dominance of the NRG1 response and the greater effect on gene downregulation was not expected, and further studies of the transcriptional regulators affected by NRG1 signalling are needed. KEGG analysis provided an indication of the type of pathways that are affected by the different ligands, but further study of these pathways and the significance of, for example, downregulation of genes linked to AMPK/mTOR by NRG1 are clearly warranted. Whilst phosphorylation of different transcription factors by ERK1/2 *vs* Akt can be expected to promote different patterns of gene expression, regulation of gene expression even just via the ERK1/2 pathway has potential to elicit different profiles. This can be effected by, for example, the degree to which ERK1/2 are activated in the nucleus, if they phosphorylate/activate downstream kinases (e.g. p90 ribosomal S6 kinases, RSKs [28]) that also phosphorylate transcription factors, the availability of different transcription factors, the affinities of kinases for those transcription factors and the affinities of transcription factors for specific gene promoters. Thus, the degree, duration and localisation of the signal can influence the genes which are expressed.

Immediate early genes include intercellular signalling components (e.g. EGFR-Ls), and transcriptional regulators (e.g. Jun, Fos, Atf3, Myc). Intercellular signalling components initiate further autocrine/paracrine signalling events with signalling cascades operating with an ever-changing panel of transcriptional regulators. As further waves of signalling/gene expression are propagated, this can lead to sustained cardiomyocyte hypertrophy, production of extracellular matrix and fibrosis or proliferation of endothelial cells and angiogenesis, features of adaptive hypertrophy. A key feature of all intracellular signalling is a need for checkpoints and systems to turn off the signal, but on-going stresses or even escalation of the problems have the potential to lead to heart failure. Thus, patterns of gene expression change with disease progression. Notably, the data in this study implicate the EGFR network all the way through to heart failure (**Figure 8**).

### The EGFR family and implications for cancer therapy-related cardiac dysfunction (CTRCD)

We used afatinib, an irreversible small molecule inhibitor developed to treat EGFR-driven cancers [52, 53]. The goal was to determine if the EGFR network is likely to play a role in development of cardiac hypertrophy in mice over the short term (7 d), not to assess the potential cardiotoxic effects of afatinib as an anti-cancer drug (unlike our studies of the BRAF/MEK inhibitors, dabrafenib and trametinib which were designed to address that issue specifically [64, 65]). Hence, we selected a high concentration designed to inhibit all EGFR family receptors rather than using a dose selective for EGFRs (the purpose of the therapy). Even so, the data have implications for cancer therapy-related cardiac dysfunction (CTRCD) resulting from therapies that target the EGFR family in that they demonstrate that inhibiting the EGFR network in a context of cardiac adaptation to pathophysiological stress affects that response. Thus, afatinib reduced the increase in LV mass induced by AngII in mice over 7 d. However, whilst there was almost complete inhibition of cardiomyocyte hypertrophy, cardiac fibrosis was increased (**Figures 5-7**). This contrasts with our studies of BRAF/MEK inhibitors in which both cardiomyocyte/cardiac hypertrophy and cardiac fibrosis were inhibited [64, 65]. The increase in fibrosis seen with afatinib and AngII suggests that pathways other than the EGFR network are involved. One possibility is that the lack of cardiomyocyte hypertrophy may have exacerbated the fibrotic response. As might be expected with enhanced fibrosis, there was some cardiac dysfunction in hearts of mice treated with afatinib and AngII. From global strain analysis, EF was preserved, but there were minor changes in GLS consistent with cardiac dysfunction, and some decrease in stroke volume and cardiac output (**Figure 3D**). Segmental analysis of the LV illustrated a significantly greater effect on peak displacement and strain in the mid- and basal segments rather than the apex of the heart (**Figure 4**). It remains to be established if there is a specific effect of the inhibitors in the basal and mid-regions. Nevertheless, the data demonstrate a detrimental effect of inhibiting EGFR family signalling on the heart.

Several inhibitors of EGFRs are used to treat cancer, including antibody therapies (e.g. anti-ERBB2/HER2 antibodies) or small molecule inhibitors (e.g. afatinib and tucatinib) [75]. Because they carry a risk of CTRCD or heart failure, guidelines have been developed for cardiac monitoring of cancer patients treated with these drugs [11, 76]. Management of CTRCD relies on consideration of traditional cardiovascular risk factors (e.g. hypertension, obesity, diabetes), early detection using cardiac imaging and traditional cardiac parameters (e.g. EF, GLS), and treatment with traditional therapies. Given the increasingly high selectivity/specificity of EGFR family inhibitors, associated CTRCD is almost certainly an on-target effect and understanding how the EGFR family network operates in cardiac cells and how anti-cancer drugs affect cardiac function (including any localised effects around the LV) may identify other options for monitoring and treatment.

### Neuregulins and cardioprotection

Cardiotoxicity associated with anti-ERBB2/HER2 antibodies used for HER2+ breast cancer (e.g. trastuzumab [77]) led to the identification of the neuregulin→ERBB2/ERBB4 network as an important cardioprotective system and the concept that exogenous administration of NRG1 (as the predominant neuregulin in the circulation) may protect the heart against pathophysiological stresses [39]. It is presumed that NRG1 activation of intracellular signalling is mediated via ERBB2/ERBB4 heterodimers but, given the receptor profiles of cardiac cells (**Figure 1B**), NRG1 could activate ERBB4 homodimers or ERBB2/ERBB4 heterodimers. Similarly, EGF could activate EGFR homodimers, or EGFR heterodimers with ERBB2, ERBB3 or ERBB4. Our data suggest that ERBB2 plays a significant and substantial role in the effects of NRG1 on gene expression, but plays a minor role in the response to EGF. Nevertheless, some of the response to EGF does appear to require ERBB2 and some of the response to NRG1 does not. This raises the question of where and how EGF family receptors operate in cardiomyocytes. The downregulation of *Nrg1* mRNA in hearts from mice treated with AngII with afatinib (**Figure 5F**) is an intriguing observation. We have observed downregulation of *Nrg1* mRNA in another mouse model with PKN1/3 double knockout further suggesting that it subject to regulation in the heart (unpublished data). Assuming *Nrg1* is required for cardiac homeostasis and Nrg1 is downregulated in disease, restoration of this signal may help to restore the homeostatic state, in which case therapies akin to recombinant rhNRG1 [39] are, indeed, likely to be beneficial for the heart, at least in some situations.

### Summary

This study demonstrates the importance of the EGFR network for the heart: in humans, the mRNA for the EGFR family and EGFR-Ls were differentially expressed in cardiac cells and expression of EGFR-L mRNAs was modulated in heart failure, EGFR-Ls were modulated in cardiomyocyte hypertrophy with differential effects on cardiomyocyte signalling/gene expression, and inhibition of EGFRs in a mouse model of hypertrophy reduced cardiac/cardiomyocyte hypertrophy, enhanced fibrosis and caused cardiac dysfunction. The data represent further steps along the way to understanding how the EGFR network operates in the heart in health and disease. Such understanding is essential to identify novel therapeutic options for cardiac disease and to develop approaches for avoiding cardiac dysfunction associated with anti-cancer drugs that target the pathways.

## Data Availability

RNASeq data are available from ArrayExpress (E-MTAB-15921) and can be accessed at https://www.ebi.ac.uk/biostudies/ArrayExpress/studies/E-MTAB-15921?key=d97f807e-4628-49f5-b6b2-12cd4334b3d9. All data are available from the authors on request.

## Abbreviations

AngII: angiotensin II;
DEG: differentially expressed gene;
DMEM: Dulbecco’s modified Eagles medium;
EF: ejection fraction;
EGF: epidermal growth factor;
EGFR: EGF receptor; EGFR-Ls,
EGFR: family ligands;
ERK: extracellular signal-regulated kinase;
ET-1: endothelin-1;
FCS: foetal calf serum;
FPKM: Fragments Per Kilobase of transcript sequence per Millions base pairs sequenced;
GLS: global longitudinal strain;
H&E: haematoxylin and eosin;
HPA: Human Protein Atlas;
LV: left ventricle;
NRG: neuregulin;
PSR: picrosirius red

