## Supplementary Tables and Figures for "The epidermal growth factor (EGF) receptor family network in cardiac hypertrophy and heart failure"

**Supplementary files in this document**

**Supplementary Table S1. Antibodies used for immunoblotting.**

**Supplementary Table S2. qPCR primers**

**Supplementary Table S3. Mouse body weights.**

**Supplementary Table S4. Single cell mRNA data for expression of EGF receptor family members and endothelin receptors in heart muscle (Human Protein Atlas version 25.0).**

**Supplementary Table S5. Single cell mRNA data for expression of EGF receptor family members and endothelin receptors in heart muscle (Human Protein Atlas version 25.0).**

**Supplementary Table S6. Echocardiography data: speckle-tracking (strain) analysis.**

**Supplementary Table S7. Single cell mRNA data for expression of EGF receptor family members and endothelin receptors in heart muscle (Human Protein Atlas version 23.0).**

**Supplementary Table S8. Single cell mRNA data for expression of EGF receptor family members and endothelin receptors in heart muscle (Human Protein Atlas version 23.0).**

**Supplementary Figure S1. Concentration-dependent effects of EGF receptor family ligands on activation of ERK1/2 or Akt in cardiomyocytes.**

**Supplementary Figure S2. Inhibition of ERBB2 with tucatinib differentially affects the changes in the cardiomyocyte transcriptome induced by EGF *vs* NRG1.**

**Supplementary Figure S3. RNASeq data for immediate early genes or EGF receptor family ligands.**

**Supplementary Figure S4. Effects of AngII ± afatinib on expression of mRNAs for fibrosis genes in mouse hearts.**

**Supplementary source File SF1. Images for Figure 2.**

**Supplementary source File SF2. Images for Figure 3A.**

**Supplementary source File SF3. Images for Figure 7A and C.**

**Supplementary source File SF4. Images for Figure 5D.**

**Supplementary Table S1. Antibodies used for immunoblotting.** All primary antibodies were from Cell Signaling Technologies Inc. Secondary antibodies were from Dako (supplied by Agilent). pAb, polyclonal antibody; mAb, monoclonal antibody.

| **Protein** | **Cat. No.** | **Host** | **Dilution** |
|---|---|---|---|
| **Primary antibodies (time course studies)** | | | |
| Phospho-Akt(Thr308) | 9275 | Rabbit pAb | 1/1000 |
| Phospho-Akt(Ser473) | 9271 | Rabbit pAb | 1/1000 |
| Total Akt | 9272 | Rabbit pAb | 1/1000 |
| Phospho-ERK1/2(T202/Y204) | 4377 | Rabbit pAb | 1/1000 |
| Total ERK1/2 | 1902 | Rabbit pAb | 1/1000 |
| **Primary antibodies (inhibitor studies)** | | | |
| Phospho-Akt(Thr308) (C31E5E) | 2905 | Rabbit mAb | 1/1000 |
| Total Akt (C67E7) | 4691 | Rabbit mAb | 1/1000 |
| Phospho-ERK1/2(T202/Y204) (D13.14.4E) | 4370 | Rabbit mAb | 1/1000 |
| Total ERK1/2 (137F5) | 4695 | Rabbit mAb | 1/1000 |
| **Secondary antibodies** | | | |
| Anti-rabbit immunoglobulins/HRP | P0448 | Goat pAb | 1/5000 |

**Supplementary Table S2. qPCR primers**

**A,** Human qPCR primers

| **Gene Symbol** | **Gene ID** | **Forward primer** | **Reverse primer** |
|---|---|---|---|
| AREG | ENSG00000109321 | GGGAAAAGTCCAYGAAAACTCACGC | GCATGTACATTTCCATTCTCTTG |
| EDNRA | ENSG00000151617 | TGGTGTGCACTGCGATCTTCTACA | ACGGCTTAAGTGAAGAGGGAACCA |
| EDNRB | ENSG00000136160 | GAAAGCCTCCGTGGGAATC | ACAGCTCGATATCTGTCAATACTCAGA |
| EGF | ENSG00000138798 | GGAAGCAATTCTCTTATTTGCTCC | GCACTACTTTCAGTTCACCAAGTGG |
| EGFR | ENSG00000146648 | GCGTCTCTTGCCGGAATGT | GGCTCACCCTCCAGAAGGTT |
| ERBB2 | ENSG00000141736 | AGCCTCTGCATTTAGGGATTCTC | CTAGCGCCGGGACGC |
| ERBB3 | ENSG00000065361 | GGTGCTGGGCTTGCTTTT | CGTGGCTGGAGTTGGTGTTA |
| ERBB4 | ENSG00000178568 | CCTGGAAGAAAGACGACTCGTTC | CGTCACTCTGATGGGTGAATTTCC |
| EREG | ENSG00000124882 | GCACAGCTTTAGTTCAGACAG | GGTCAAAGCCACATATTCTTTGC |
| GAPDH | ENSG00000111640 | CCAAGGTCATCCATGACAACTT | AGGGGCCATCCACAGTCTT |
| HBEGF | ENSG00000113070 | CGGAAAGTTCCGTGACTTGCAAGAG | CCTCTCTCCATGGTAACCCGGCTG |
| NRG1 | ENSG00000157168 | CAAGTGGTTCAAGAATGGGA | GAAGTTCTGACTTCCCTGG |
| TGFA | ENSG00000163235 | CCAGATCCCACACTCAGTTCTGC | GGACCTGGCAGCAGTGTATCAGC |

**B,** Mouse qPCR primers

| Gene Symbol | Gene ID | Forward primer | Reverse primer |
|---|---|---|---|
| Areg | ENSMUSG00000029378 | CGCTTATGGTGGAAACCTCTC | GGTCTTAGGCTCAGGCCATTA |
| Ccn2 (Ctgf) | ENSMUSG00000019997 | GCACACCGCACAGAACCA | ATGGCAGGCACAGGTCTTG |
| Col1a1 | ENSMUSG00000001506 | TCGTGGCTTCTCTGGTCTC | CCGTTGAGTCCGTCTTTGC |
| Col2a1 | ENSMUSG00000022483 | GACGAGGCAGACAGTACCTTG | GATGCTCTCAATCTGGTTGTTCAG |
| Col3a1 | ENSMUSG00000026043 | GGAACCTGGTTTCTTCTCACC | TAGGACTGACCAAGGTGGCT |
| Col4a1 | ENSMUSG00000031502 | TGTGGGCCAGCCAGGCATTG | CAGGGGGTCCGATCGCTCCA |
| Egf | ENSMUSG00000028017 | AGCATACTCAGCGTCACAGC | GCAGGACCGGCACAAGTC |
| Ereg | ENSMUSG00000029377 | CACCGAGAAAGAAGGATGGAG | TGTGCTGATAACTGCCTGTAG |
| Fn1 | ENSMUSG00000026193 | AAGAGGACGTTGCAGAGCTA | AGACACTGGAGACACTGACTAA |
| Gapdh | ENSMUSG00000057666 | TCACCACCATGGAGAAGGC | GCTAAGCAGTTGGTGGTGCA |
| Hbegf | ENSMUSG00000024486 | CGGGGAGTGCAGATACCTG | TTCTCCACTGGTAGAGTCAGC |
| Myh7 | ENSMUSG00000053093 | CATGCCAACCGTATGGCTG | GTTCCACGATGGCGATGTTC |
| Nppa | ENSMUSG00000041616 | GATGGATTTCAAGAACCTGCTAGA | CTTCCTCAGTCTGCTCACTCA |
| Nppb | ENSMUSG00000029019 | TCCAGCAGAGACCTCAAAATTC | CAGTGCGTTACAGCCCAAA |
| Nrg1 | ENSMUSG00000062991 | GAGAACACCCAAGTCAGGA | GATGTAGATGTGGAAAGTTTAGGAG |
| Postn | ENSMUSG00000027750 | TTCCTCTCCTGCCCTTATATGC | CCTGATCCCGACCCCTGAT |
| Tagln (Sm22-a) | ENSMUSG00000032085 | GACTGCACTTCTCGGCTCAT | CCGAAGCTACTCTCCTTCCA |
| Tgfa | ENSMUSG00000029999 | CTCTGCTAGCGCTGGGTATC | TGGGCACTTGTTGAAGTGAG |
| Tgfb1 | ENSMUSG00000002603 | TGGACACACAGTACAGCAAG | GTAGTAGACGATGGGCAGTG |

**C,** Rat qPCR primers

| Gene Symbol | Gene ID | Forward primer | Reverse primer |
|---|---|---|---|
| Agtr1a | ENSRNOG00000018346 | GATCACCAGGTCAAGTGGA | GCAGAAGAGTTAAGGGCCA |
| Areg | ENSRNOG00000002754 | TCGCAGCTATTGGCATCATTA | TTCTGCTTCTTCATATTCCCTGAA |
| Atf3 | ENSRNOG00000003745 | AGCAGGATCGCACTAATGGG | ACAACTTCAATGATGATGAATGTTCTCAC |
| Ddr2 | ENSRNOG00000002881 | GGAAACAGGTGCTTGATGG | CAAATCTAGCTACAATGGGTGG |
| Dusp6 | ENSRNOG00055005953 | AGCGACTGGAATGAGAACACA | GCAGCCCTCGTCTTTGAGT |
| Egf | ENSRNOG00000053979 | TGCCAACTGGGGGTGCACAG | CTGCCCGTGGCCAGCGTGGC |
| Egfr | ENSRNOG00000004332 | CCACCAAGACAGGCGACG | AGTAGCTTGGTTCTCGCAGTC |
| Egr1 | ENSRNOG00000019422 | TCAGTCGTAGTGACCACCTTAC | GGTATGCCTCTTGCGTTCATC |
| Erbb2 | ENSRNOG00000006450 | CACTGTACCTTGGGACCAG | CCTCGAGACCACAATCCTC |
| Erbb3 | ENSRNOG00000004964 | CATAAGATACCCGCACTGGA | CTGGATGTTTAGGTAACCTGTG |
| Erbb4 | ENSRNOG00000014248 | CTCCAATAGGAGTGAAATTGGAC | CCATCCTGGTACACAAACTG |
| Ereg | ENSRNOG00000002771 | GGAAGAATACGAGAGAGTGACCTC | GCTGCACATCCTTGTCCACA |
| Fos | ENSRNOG00000008015 | AGTAGAGCAGCTATCTCCTG | AGTTGATCTGTCTCCGCTTG |
| Gapdh | ENSRNOG00000018630 | CCAAGGTCATCCATGACAACTT | AGGGGCCATCCACAGTCTT |
| Hbegf | ENSRNOG00000018646 | GGAGAGTGCAGATACCTGAAGGA | GTCAGCCCATGACACCTCTGT |
| Jun | ENSRNOG00000026293 | GATCATCCAGTCCAGCAATG | TATTCTGGCTATGCAGTTCAG |
| Mhy7 | ENSRNOG00000016983 | TCACCTACCAGACAGAGG | CCTTCACCTTTAACTGCAGC |
| Myc | ENSRNOG00000004500 | CGTTATTTGAAGCCTGAATTTCCT | TCATAGTTCCTGTTAGCGAAGC |
| Myh6 | ENSRNOG00000025757 | TACCAGAAGTCCTCCCTCAAAC | CTTGCCTCCTTTGCCTTTCC |
| Mhy7 | ENSRNOG00000016983 | TCACCTACCAGACAGAGG | CCTTCACCTTTAACTGCAGC |
| Nppb | ENSRNOG00000008141 | CCAGGAGAGACTTCGAAATTCCA | TGCAGCCAGGAGGTCTTCC |
| Nrg1 | ENSRNOG00000010392 | CTACCAGAAGAGGGTGCTGACAA | GCCGCTGCTTCTTGGTTTT |
| Tagln (Sm22a) | ENSRNOG00000017628 | GTTTATGAAGAAAGCCCAGGA | TTTGAAGGCCAATGACGTG |
| Tgfa | ENSRNOG00000016182 | ATGGTCCCCTCGGCTGGA | GCTGCTTCTTCTGGCTGGCA |
| Tie2/Tek | ENSRNOG00000002881 | GAGGTTCTTTGTAAACTTGGAC | CAGGTATAAGTAGCCTCTGTG |

**Supplementary Table S3. Mouse body weights.**

| | Starting body weight (g) | | Body weight after MP surgery (g) | | End body weight (g) | |
|---|---|---|---|---|---|---|
| | **Mean** | **SD** | **Mean** | **SD** | **Mean** | **SD** |
| **3 d qPCR study** | | | | | | |
| Vehicle (n=5) | --- | --- | 24.6 | 1.1 | 24.9 | 1.2 |
| AngII (n=4) | --- | --- | 23.8 | 1.2 | 24.0 | 1.3 |
| **Afatinib/AngII study** | | | | | | |
| Vehicle (n=7) | 24.2 | 2.4 | 26.1 | 2.8 | 26.7 | 2.8 |
| AngII (n=7) | 24.2 | 1.8 | 26.6 | 1.7 | 26.9 | 2.0 |
| Afatinib/AngII (n=7) | 24.2 | 1.3 | 26.4 | 1.1 | 26.9 | 1.6 |

**Supplementary Table S4. Single cell mRNA data for expression of EGF receptor family members and endothelin receptors in heart muscle (Human Protein Atlas version 25.0).** Normalised data are provided (nCPM).

| | **Cell type** | **EGFR** | **ERBB2** | **ERBB3** | **ERBB4** | **EDNRA** | **EDNRB** |
|---|---|---|---|---|---|---|---|
| c-0 | Cardiomyocytes | 9.5 | 104.4 | 0.6 | 1157.3 | 82.7 | 3.3 |
| c-1 | Cardiomyocytes | 18.5 | 98.6 | 1.4 | 900.1 | 83.8 | 4.2 |
| c-2 | Cardiomyocytes | 13.6 | 93.2 | 1.1 | 874.7 | 86.2 | 4 |
| c-3 | Cardiomyocytes | 15 | 126.4 | 0.9 | 932.9 | 72.4 | 4.1 |
| c-5 | Cardiomyocytes | 13.4 | 83.7 | 1.5 | 876.2 | 114.5 | 5.4 |
| c-9 | Cardiomyocytes | 5.8 | 59.9 | 0 | 1046.8 | 61.1 | 2.3 |
| c-10 | Cardiomyocytes | 13.4 | 77.5 | 0 | 916.7 | 68.2 | 0 |
| c-12 | Cardiomyocytes | 2.6 | 139.4 | 0 | 465.5 | 31.6 | 0 |
| c-4 | Fibroblasts | 392.2 | 73.8 | 1.9 | 36.5 | 11 | 90.5 |
| c-11 | Epicardial cells | 477.6 | 40.9 | 2.3 | 209.2 | 22.7 | 11.4 |
| c-14 | Adipocytes | 193.3 | 17.8 | 0 | 15.3 | 22.9 | 10.2 |
| c-7 | Endothelial cells | 62.2 | 86.7 | 1.4 | 29.4 | 4.2 | 68.5 |
| c-8 | Pericytes | 159.7 | 55.2 | 1.5 | 20.9 | 302.9 | 107.4 |
| c-17 | Pericytes | 49.4 | 39.5 | 0 | 19.8 | 168.1 | 138.4 |
| c-13 | Smooth muscle cells | 85.1 | 71 | 3.5 | 145.5 | 259 | 35.5 |
| c-6 | Macrophages | 16.9 | 54.9 | 0 | 73.2 | 8 | 8.4 |
| c-15 | T-cells | 103.7 | 95.7 | 0 | 27.9 | 0 | 51.8 |
| c-16 | Mast cells | 117.4 | 29.4 | 0 | 36.7 | 14.7 | 80.7 |

**Supplementary Table S5. Single cell mRNA data for expression of EGF receptor family members and endothelin receptors in heart muscle (Human Protein Atlas version 25.0).** Normalised data are provided (nCPM). CMs, cardiomyocytes

| | Cell type | EGF | AREG | EREG | HB-EGF | TGFA | NRG1 | NRG2 | NRG3 | NRG4 | EDN1 |
|---|---|---|---|---|---|---|---|---|---|---|---|
| c-0 | CMs | 69.2 | 4.6 | 10.5 | 3.9 | 6.9 | 3.5 | 113.4 | 5.8 | 40.4 | 0.8 |
| c-1 | CMs | 45.1 | 3 | 12.8 | 2.9 | 6.2 | 5.7 | 95.8 | 10.7 | 72.9 | 0.8 |
| c-2 | CMs | 71.2 | 3.5 | 4.6 | 4.6 | 9.7 | 6 | 104.8 | 10.9 | 51.9 | 0.5 |
| c-3 | CMs | 49.8 | 3.8 | 10.7 | 4.6 | 7.3 | 4.7 | 94.4 | 18.3 | 49.2 | 0.5 |
| c-5 | CMs | 56.6 | 6.6 | 9.8 | 6.1 | 7.1 | 6.8 | 136.2 | 10 | 44.7 | 0.7 |
| c-9 | CMs | 49.6 | 20.8 | 8.1 | 19.6 | 11.5 | 4.6 | 88.8 | 11.5 | 45 | 1.2 |
| c-10 | CMs | 66.8 | 1.3 | 5.3 | 5.3 | 20 | 8 | 80.2 | 12 | 45.4 | 1.3 |
| c-12 | CMs | 26.3 | 0 | 0 | 5.3 | 0 | 2.6 | 97.3 | 7.9 | 71 | 0 |
| c-4 | Fibroblasts | 1 | 0.6 | 1 | 3.7 | 1 | 13.3 | 10.2 | 16.5 | 70.6 | 2.1 |
| c-11 | Epicardial cells | 0 | 232 | 9.1 | 11.4 | 2.3 | 11.4 | 22.7 | 11.4 | 68.2 | 11.4 |
| c-14 | Adipocytes | 5.1 | 2.5 | 0 | 0 | 0 | 7.6 | 38.2 | 10.2 | 68.7 | 0 |
| c-7 | Endothelial cells | 4.2 | 3.5 | 0.7 | 2.8 | 0 | 326.4 | 9.1 | 698.9 | 71.3 | 14.7 |
| c-8 | Pericytes | 0 | 1.5 | 0 | 3 | 0 | 9 | 9 | 4.5 | 22.4 | 1.5 |
| c-17 | Pericytes | 0 | 0 | 0 | 0 | 0 | 0 | 9.9 | 0 | 49.4 | 0 |
| c-13 | Smooth muscle cells | 0 | 0 | 3.5 | 3.5 | 0 | 0 | 10.6 | 14.2 | 67.4 | 0 |
| c-6 | Macrophage | 2.3 | 13.1 | 3.8 | 13.6 | 11.7 | 7 | 4.2 | 9.4 | 59.1 | 0 |
| c-15 | T-cells | 0 | 0 | 0 | 4 | 8 | 8 | 8 | 27.9 | 47.8 | 0 |
| c-16 | Mast cells | 0 | 44 | 0 | 0 | 0 | 29.4 | 7.3 | 7.3 | 14.7 | 0 |

**Supplementary Table S6. Echocardiography data: speckle-tracking (strain) analysis.** Male C57Bl/6J mice (9 weeks) received a baseline echocardiogram with subsequent echocardiograms taken 3 d and 7 d after osmotic minipumps were implanted to deliver vehicle, 0.8 mg/kg/d AngII or AngII with 4.5 mg/kg/d afatinib (n=7 per group). EDV, end diastolic volume; ESV, end systolic volume; EDLVM, end diastolic LV mass; ESLVM, end systolic LV mass; GLS, global longitudinal strain; GCS, global circumferential strain.

| | Vehicle | | AngII | | Afatinib/AngII | |
|---|---|---|---|---|---|---|
| | **Mean** | **SD** | **Mean** | **SD** | **Mean** | **SD** |
| **Baseline** | | | | | | |
| Heart Rate (bpm) | 466 | 16 | 460 | 31 | 452 | 40 |
| Stroke volume (µl) | 28.49 | 4.40 | 35.37 | 10.18 | 28.61 | 7.24 |
| Ejection fraction (%) | 50.23 | 4.63 | 55.61 | 5.89 | 49.80 | 8.61 |
| Fractional shortening (%) | 25.39 | 3.46 | 29.44 | 5.31 | 25.91 | 5.63 |
| Cardiac output (ml/min) | 13.30 | 2.22 | 16.20 | 4.65 | 12.94 | 3.58 |
| EDV (µl) | 57.33 | 12.69 | 63.07 | 16.79 | 56.82 | 6.57 |
| ESV (µl) | 28.84 | 8.63 | 27.70 | 8.08 | 28.22 | 4.80 |
| EDLVM (mg) | 54.53 | 8.31 | 58.16 | 6.84 | 54.39 | 4.18 |
| ESLVM (mg) | 56.71 | 9.27 | 62.40 | 9.37 | 56.21 | 4.31 |
| GLS (%) | -17.97 | 2.71 | -19.95 | 3.43 | -17.70 | 4.07 |
| GCS (%) | -17.76 | 2.77 | -19.01 | 2.39 | -19.86 | 4.27 |
| **3 d** | | | | | | |
| Heart Rate (bpm) | 474 | 39 | 492 | 48 | 498 | 56 |
| Stroke volume (µl) | 31.99 | 8.62 | 29.48 | 8.59 | 23.75 | 6.77 |
| Ejection fraction (%) | 56.59 | 5.56 | 58.95 | 7.21 | 55.10 | 2.51 |
| Fractional shortening (%) | 30.66 | 5.29 | 29.93 | 7.99 | 30.51 | 4.81 |
| Cardiac output (ml/min) | 15.16 | 3.96 | 14.44 | 4.12 | 11.93 | 4.08 |
| EDV (µl) | 56.01 | 12.35 | 50.84 | 18.27 | 42.97 | 12.36 |
| ESV (µl) | 24.02 | 5.17 | 21.36 | 10.78 | 19.22 | 5.78 |
| EDLVM (mg) | 55.21 | 7.84 | 69.00 | 7.46 | 60.74 | 5.13 |
| ESLVM (mg) | 57.48 | 7.80 | 71.94 | 9.49 | 62.13 | 5.96 |
| GLS (%) | -19.61 | 1.53 | -21.37 | 3.89 | -16.80 | 2.61 |
| GCS (%) | -21.70 | 4.13 | -21.07 | 2.88 | -18.92 | 4.94 |
| **7 d** | | | | | | |
| Heart Rate (bpm) | 504 | 62 | 514 | 43 | 493 | 74 |
| Stroke volume (µl) | 30.91 | 6.28 | 27.94 | 6.25 | 23.99 | 4.72 |
| Ejection fraction (%) | 59.49 | 6.09 | 57.70 | 8.56 | 56.29 | 8.69 |
| Fractional shortening (%) | 32.54 | 6.82 | 31.86 | 8.24 | 29.49 | 7.79 |
| Cardiac output (ml/min) | 15.58 | 3.66 | 14.21 | 2.54 | 11.66 | 2.15 |
| EDV (µl) | 52.17 | 12.08 | 49.91 | 17.09 | 43.65 | 11.81 |
| ESV (µl) | 21.26 | 6.71 | 21.97 | 12.38 | 19.66 | 8.71 |
| EDLVM (mg) | 54.48 | 7.81 | 74.82 | 7.08 | 65.62 | 3.08 |
| ESLVM (mg) | 56.42 | 7.94 | 77.33 | 5.70 | 66.87 | 2.62 |
| GLS (%) | -20.49 | 2.53 | -19.38 | 3.53 | -17.87 | 3.35 |
| GCS (%) | -23.18 | 2.83 | -20.90 | 2.41 | -23.50 | 3.03 |

**Supplementary Table S7. Single cell mRNA data for expression of EGF receptor family members and endothelin receptors in heart muscle (Human Protein Atlas version 23.0).** Normalised data are provided (nCPM).

| | Cell type | EGFR | ERBB2 | ERBB3 | ERBB4 | EDNRA | EDNRB |
|---|---|---|---|---|---|---|---|
| c-0 | Cardiomyocytes | 1.8 | 24 | 0.2 | 16.3 | 24.3 | 4.6 |
| c-1 | Cardiomyocytes | 5.7 | 20.1 | 2.4 | 27.7 | 7.6 | 2.4 |
| c-2 | Cardiomyocytes | 15.4 | 61.7 | 0.4 | 48.3 | 47.8 | 21 |
| c-5 | Cardiomyocytes | 8.7 | 56.3 | 1.4 | 38.5 | 27.9 | 8.2 |
| c-8 | Cardiomyocytes | 33.6 | 24.3 | 11.3 | 28.1 | 52.4 | 2.4 |
| c-7 | Fibroblasts | 101.3 | 16.8 | 22.8 | 3.5 | 4.2 | 77.1 |
| c-4 | Endothelial cells | 15.2 | 10.2 | 0.4 | 0 | 2.7 | 179.3 |
| c-6 | Endothelial cells | 34.7 | 8.4 | 3.9 | 0.1 | 2.2 | 205 |
| c-3 | Smooth muscle cells | 37.3 | 11.1 | 3.8 | 1.2 | 92.4 | 137.6 |

**Supplementary Figure S1. Concentration-dependent effects of EGF receptor family ligands on activation of ERK1/2 or Akt in cardiomyocytes.** Rat neonatal cardiomyocytes were treated with the indicated concentrations of AREG, EREG, HBEGF, NRG1 or TGFA (5 min). Proteins were immunoblotted with antibodies to phosphorylated (P) or total (T) ERK1/2 (**A**) or Akt(Thr308) (**B**). Densitometric data are provided as means ± SEM (n=4 independent cardiomyocyte preparations) and are presented as the ratio of phosphorylated:total protein, normalised to the mean of the controls.

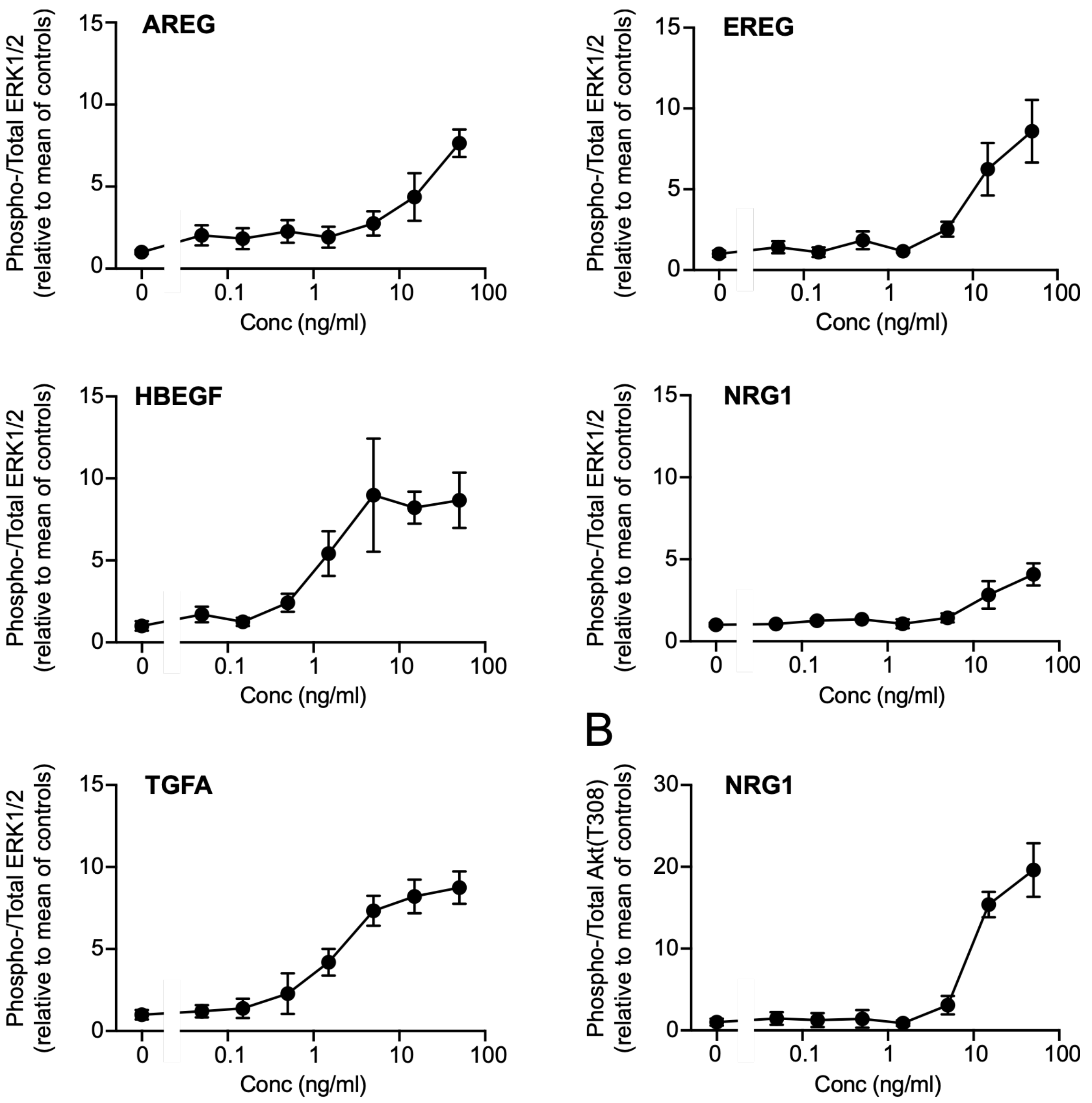

**Supplementary Figure S2. Inhibition of ERBB2 with tucatinib differentially affects the changes in the cardiomyocyte transcriptome induced by EGF *vs* NRG1.** Rat neonatal cardiomyocytes were treated with EGF or NRG1 (100 ng/ml, 60 min) in the absence or presence of 1 μM tucatinib (tuc) and mRNA expression measured by RNASeq. Differentially expressed genes (DEGs) were identified (FDR<0.05). Data show the comparison of expression levels of gene clusters significantly upregulated or downregulated by EGF (A) or NRG1 (B) and unaffected (not inhibited) or significantly inhibited by tucatinib. Data are presented as means ± SEM (n values as indicated). Statistical analysis used repeated measures one-way ANOVA with Geisser-Greenhouse correction and Tukey's multiple comparisons test.

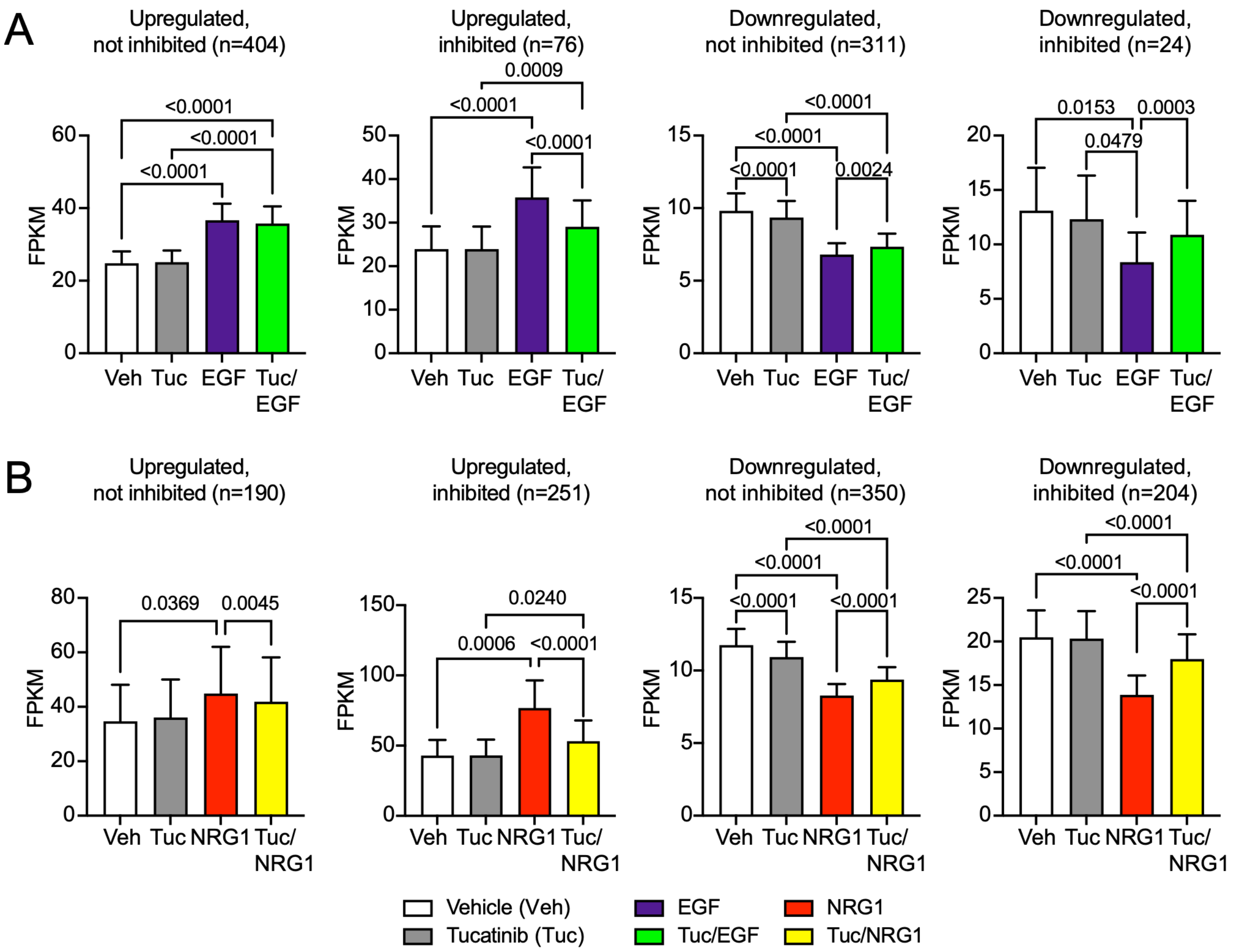

**Supplementary Figure S3. RNASeq data for immediate early genes or EGF receptor family ligands.** Rat neonatal cardiomyocytes were treated with EGF or NRG1 (100 ng/ml, 60 min) in the absence or presence of 1 µM tucatinib (tuc) and mRNA expression measured by RNA-Seq. Data are provided for selected immediate early genes (**A**) or EGF receptor family ligands (**B**) for comparison with qPCR analysis.

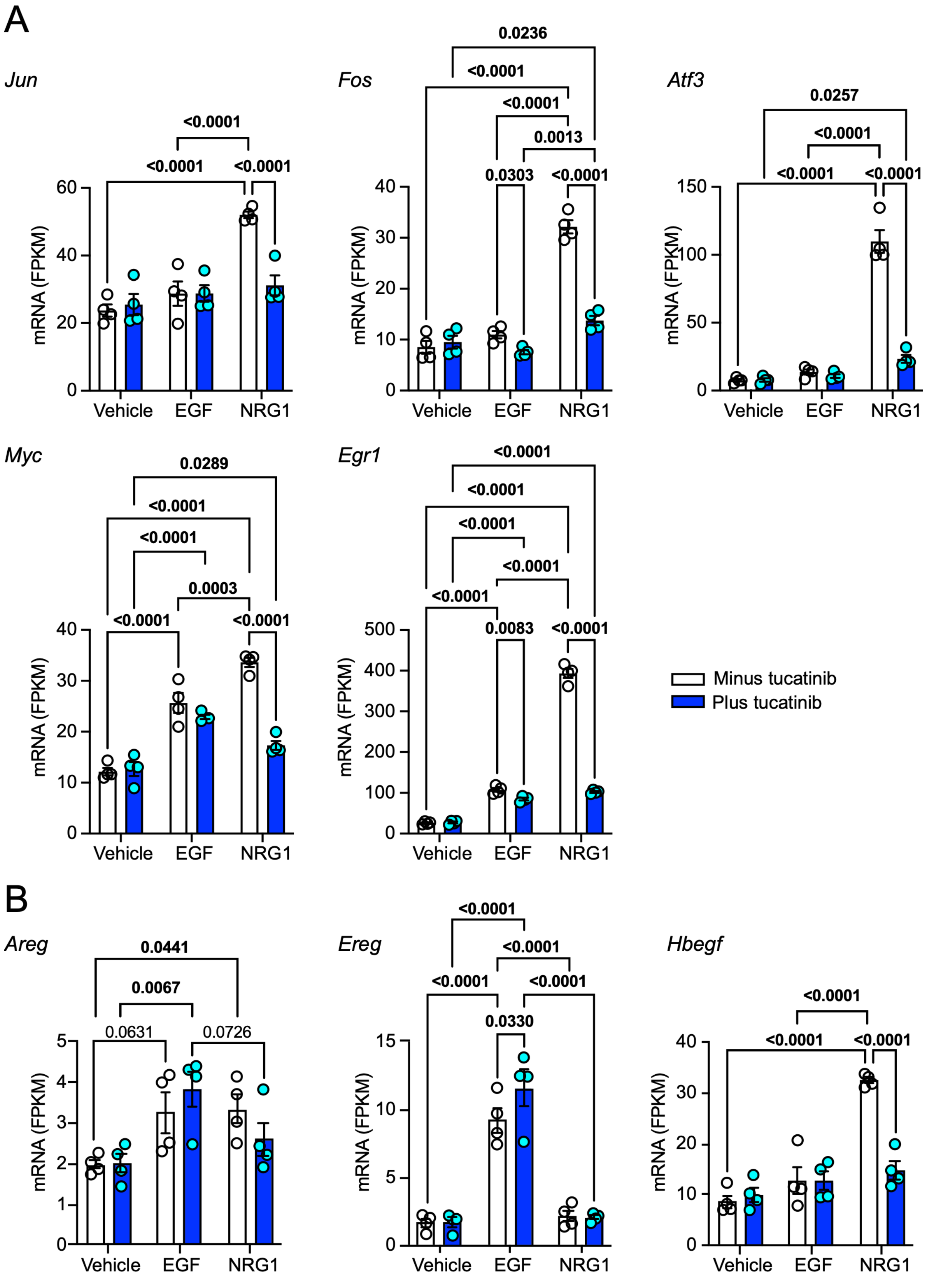

**Supplementary Figure S4. Effects of AngII ± afatinib on expression of mRNAs for fibrosis genes in mouse hearts.** Male C57Bl/6J mice were treated with vehicle, 0.8 mg/kg/d AngII or AngII with 4.5 mg/kg/d afatinib for 7 d (n=7 per group). Hearts were ground to powder under liquid $N_2$ and samples used to prepare RNA for qPCR. Statistical analysis used one-way ANOVA with Tukey's post-hoc tests. Significant (p<0.05) changes are shown in bold type.

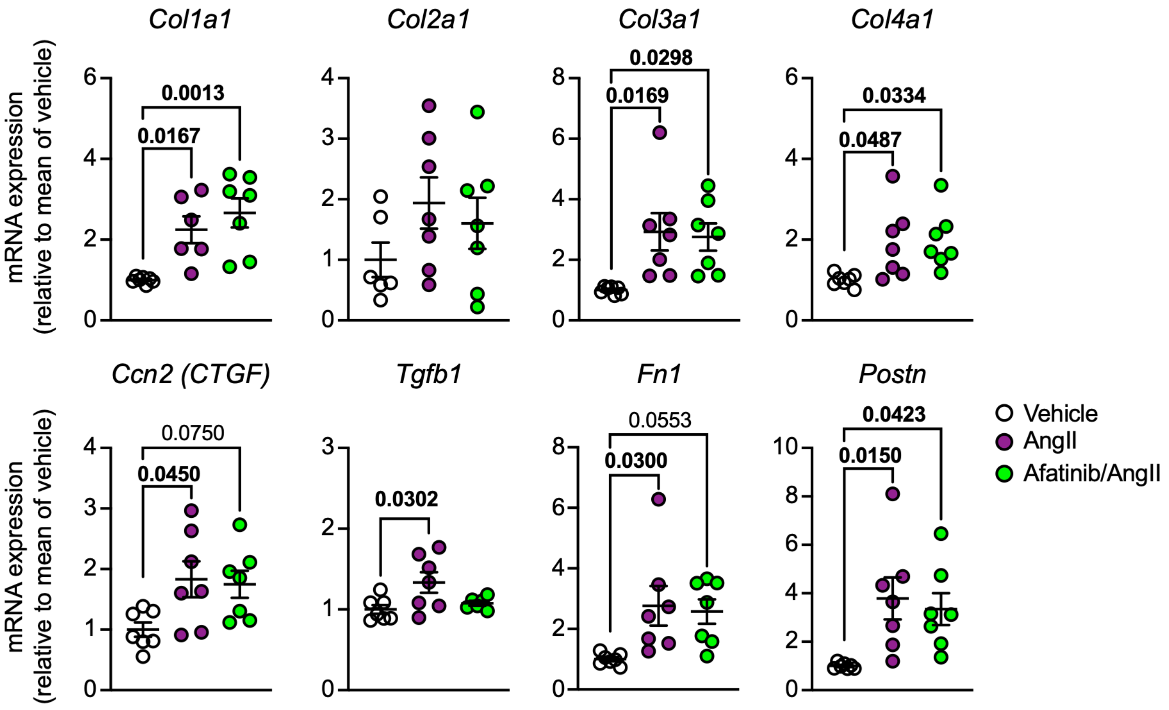

### Supplementary Source Figure SF1. Images for Figure 2

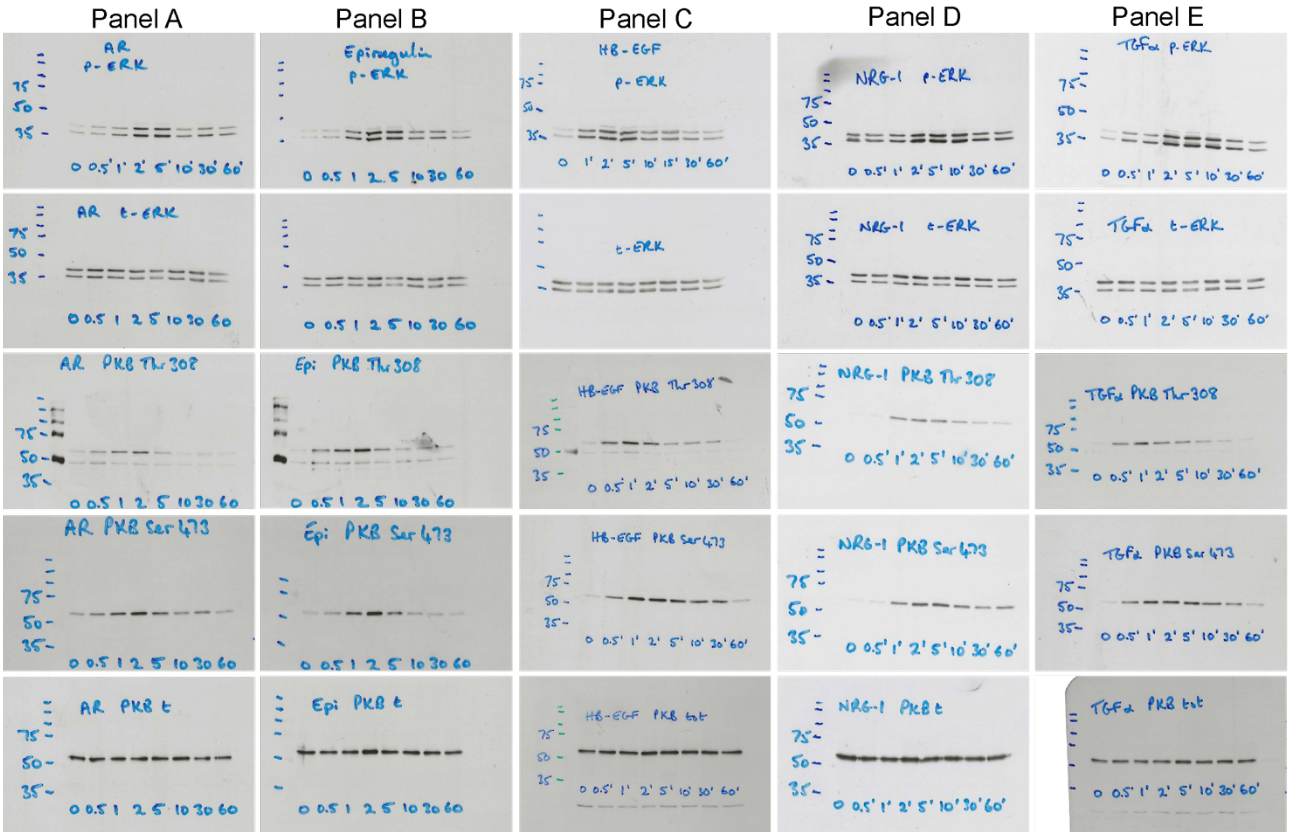

### Supplementary Source Figure SF2. Images for Figure 3A

Blot probed with antibodies to phosphorylated ERK1/2 and Akt(T308)

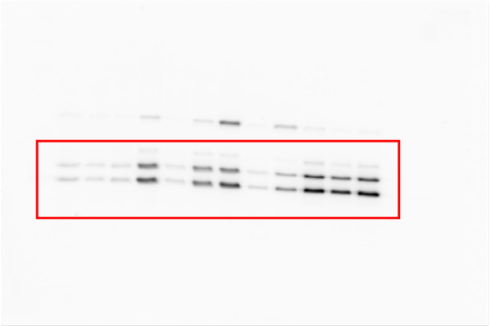

Image used for phosphorylated ERK1/2

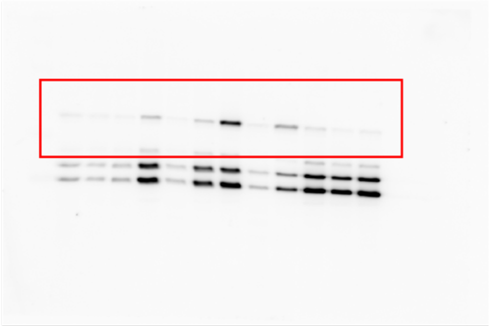

Image used for phosphorylated Akt(T308)

Blot probed with antibodies to total ERK1/2 and Akt

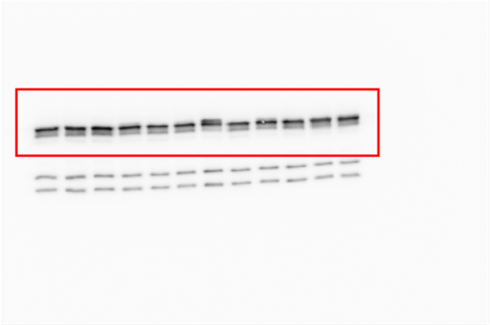

Image used for total ERK1/2

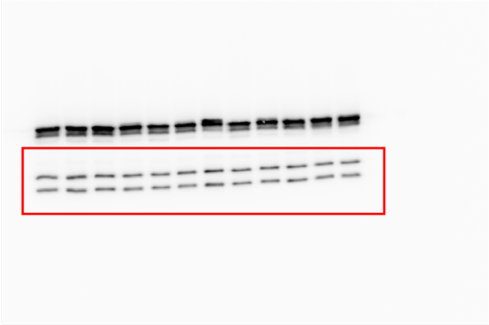

Image used for total Akt(T308)

**Supplementary Source Figure SF3. Images for Figure 7A and C.**

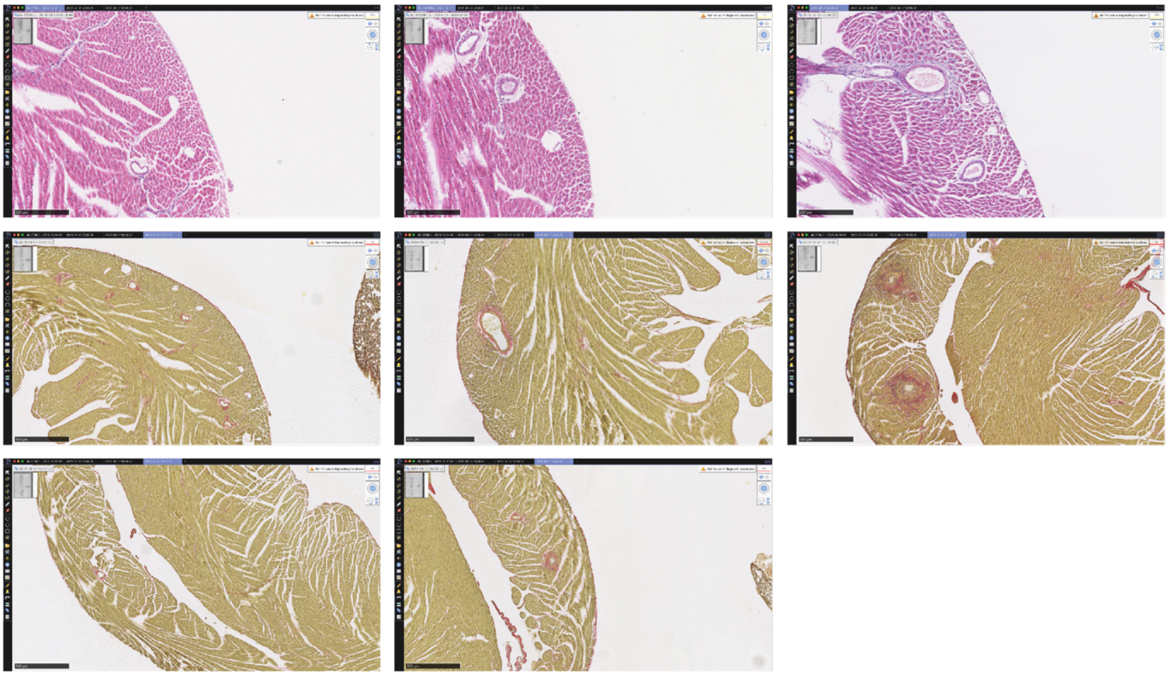

**Supplementary Source Figure SF4. Images for Figure 5D.**

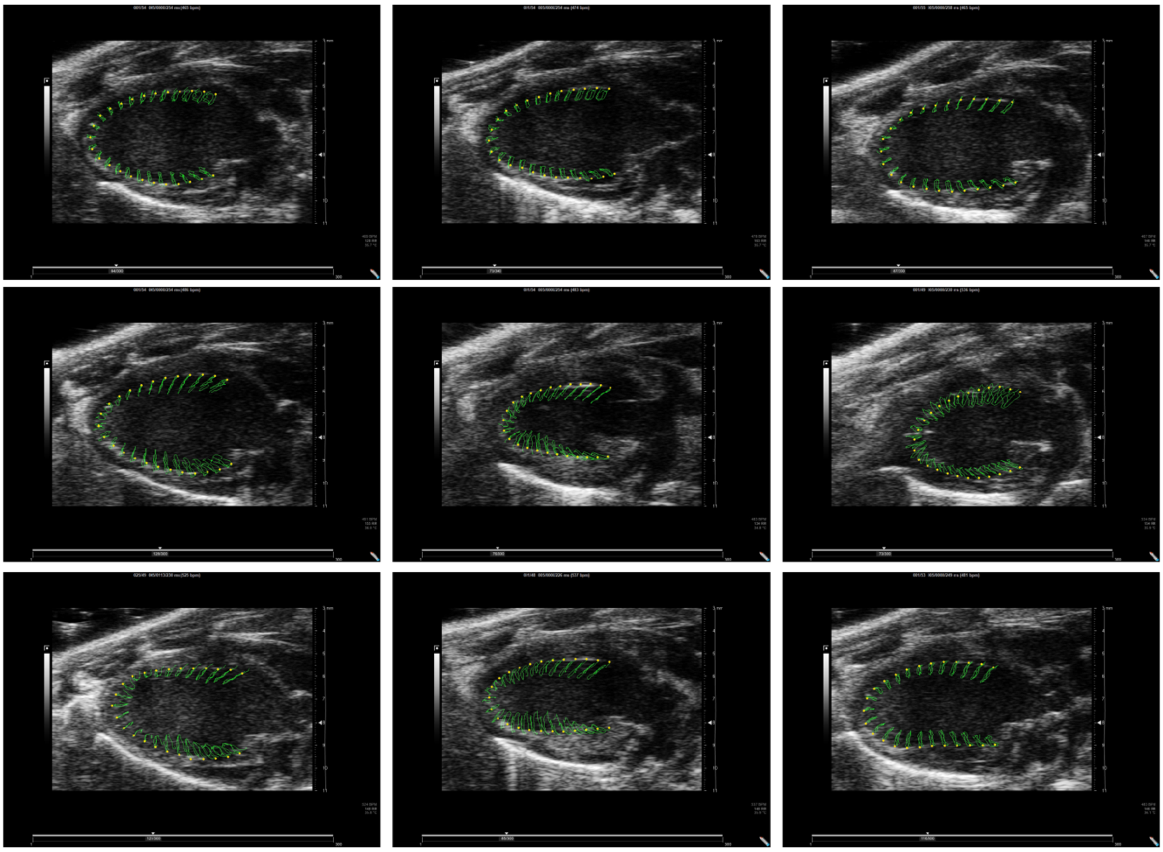
